# The sugar-beet cyst nematode effector *Hs*2B11 targets the Arabidopsis serine protease inhibitor *At*PR-6 to favor parasitism

**DOI:** 10.64898/2026.02.21.707195

**Authors:** Joffrey Mejias, Melissa Bredow, Anil Kumar, Parijat S. Juvale, Thomas R. Maier, Ekkachai Khwanbua, Steven A. Whitham, Sebastian Eves-van den Akker, Thomas J. Baum

## Abstract

Cyst nematodes secrete effector proteins to manipulate host cell biology and suppress immunity, yet the mechanisms underlying these interactions remain largely unexplored. In this study, we characterize the function of *Hs*2B11, a *Heterodera schachtii* effector that was previously shown to be expressed in the dorsal gland of sugar-beet cyst nematodes (BCN). Here, we report that in *Arabidopsis thaliana Hs2*B11 functions as an immune regulator that modulates the production of elicitor-induced oxidative species, likely to favor parasitism. To elucidate the molecular basis of this immune suppression, we performed a yeast-two-hybrid screen and identified the host serine protease inhibitor *At*PR-6 as a direct interactor of *Hs*2B11. We show that *At*PR-6 acts as a positive regulator of plant immunity; its expression is induced upon nematode infection and knock-out of *AtPR-6* compromises oxidative species production leading to higher susceptibility to *H. schachtii* infection. Conversely, *AtPR-6* overexpression enhances immune responses resulting in increased resistance to BCN infection. Detailed analysis of this interaction demonstrated that *Hs*2B11 interacts with *At*PR-6 using its carboxyl-terminal domain. AlphaFold2 predicts that the C-terminal domain forms a beta-solenoid-like structure with a ladder of serine residues organized across one of its surfaces. We propose that using this interface, *Hs*2B11 targets *At*PR-6 via molecular titration, preventing the inhibitor from regulating host proteases that control immune signaling. These findings highlight a counter-defense strategy where a nematode effector neutralizes a specific host protease inhibitor to subvert plant immunity.

## Introduction

Soybean (*Glycine max*) is one of the most important sources of food and feed production in the world [1]. Importantly, as soybean forms symbiotic relationships with nitrogen-fixing bacteria, its cultivation reduces the need for exogenous fertilizer applications, supporting sustainable agriculture practices [2]. The most destructive pathogen of soybean is the soybean cyst nematode (SCN), *Heterodera glycines*, which causes yield losses estimated to be billions of dollars each year [3]. To maintain soybean crops at their highest production level, it is crucial to identify novel resistance sources or mechanisms that can be exploited to combat this pathogen.

SCN infects soybean by delivering effector molecules to hijack the plant’s physiology, including a complete reprogramming of a selected root cell and the inactivation of plant defense mechanisms [4]. The sugar-beet cyst nematode (BCN), *H. schachtii,* is an effective model for studying cyst nematode pathogenicity, as it infects the model plant *Arabidopsis thaliana* (hereinafter, Arabidopsis). Cyst nematodes are attracted to host roots through chemoattraction and invade them to migrate into the vascular cylinder, inducing the formation of a reprogrammed feeding structure called a syncytium. The formation of this feeding structure allows nematodes to switch to a sedentary lifestyle and feed exclusively on those newly formed cell types. Through its nutrient and energy acquisition from the host, nematodes molt several times to become a female able to produce hundreds of eggs. The female will subsequently die and harden to form a “cyst” containing the eggs. Finally, new juveniles will hatch from those eggs when environmental conditions are favorable [5].

The plant immune system needs to be drastically suppressed during the entire life cycle of the nematode, including penetration and formation of the feeding site. Proteinaceous effectors are keystones of pathogenicity and essential to manipulate plant physiology [6]. BCNs produce hundreds of secreted effector proteins, primarily in the salivary glands, which are composed of two subventral gland cells and one dorsal gland cell in cyst-nematodes. The subventral glands are involved in the production of secreted cell-wall modifying enzymes that aid the migration of nematodes into the root. The dorsal gland becomes more active at parasitic stages of infection and is mostly involved in the production of effector proteins that suppress plant immunity, or initiate and maintain the syncytium. Investigating the function of effectors capable of suppressing plant defenses, and identifying their plant targets, could pinpoint key resistance genes or susceptibility genes that can be engineered to create durable resistance against cyst-nematodes [7].

Among the large range of functions associated with effector proteins, those harboring protease activity have been shown to be critical for pathogenicity. Secreted proteases possessing virulence activity have been described for phytopathogenic bacteria, fungi, viruses and nematodes [8–11], suggesting that proteases are used as common weapons to favor infection. The effector AvrPphB, from *Pseudomonas syringae,* is one the most characterized secreted protease effectors, which specifically targets a subfamily of receptor-like cytoplasmic kinases to suppress defense responses induced by cell surface-localized plant immune receptors such as FLAGELLIN SENSITIVE 2 (FLS2) [12]. Arabidopsis or soybean carrying the intracellular nucleotide-binding leucine-rich repeat (NLR) protein RPS5 (RESISTANCE TO PSEUDOMONAS SYRINGAE 5) indirectly detect AvrpPhB, by detecting the cleavage product of the Arabidopsis protein kinase AvrpPhB-Susceptible 1, leading a hypersensitive cell death response, and resistance against *P. syringae* [13]. Numerous proteases have been detected in the transcriptome or secretome of phytoparasitic nematodes, underscoring their importance during interaction with host plants [14–16]. In phytoparasitic nematodes, secreted proteases have been proposed to function in helping to digest the intracellular contents of plant cells, which are thereafter used as a nutrient source, or to hijack host proteins [14–16]. For example, the SCN effector Cysteine Protease 1 (CPR1) targets a branched-chain amino acid aminotransferase from soybean enhancing susceptibility to SCN [17].

In order to counteract the effect of secreted protease effectors, plants produce protease inhibitors, some of which have been demonstrated to localize to the extracellular space of plant cells to covalently link and inhibit pathogen secreted proteases. For example, the activity of two subtilisin-like and trypsin-like proteases from *Fusarium culmorum* are inhibited by three barley (*Hordeum vulgare*) chymotrypsin/subtilisin protease inhibitors, CI-1A, -1B, and -2 [18]. Similarly, a trypsin-chymotrypsin protease inhibitor from broad bean (*Vicia faba*) has demonstrated antifungal activity against *Mycosphaerella arachidicola*, *F. oxysporum*, and *Botrytis cinerea* [19].

Protease inhibitors also directly modulate plant immunity by fine-tuning the activity of endogenous plant proteases. For example, microbe-associated molecular pattern (MAMP)-induced oxidative species production is tightly controlled by the activity of a papain-like cysteine protease, XYLEM CYSTEINE PEPTIDASE 1 (XCP1), and its inhibitor CYSTATIN 6 (CYS6) in Arabidopsis [20]. Alternatively, pathogens can secrete protease inhibitors to target plant proteases such as the cysteine-protease inhibitor Avr2 from *Cladosporium fulvum* that binds and inhibits the tomato (*Solanum lycopersicum*) cysteine protease Rcr3pim [21]. This interaction is sensed by the extracellular immune receptor Cf-2, which is required to mount an immune response against this pathogen. Comparatively less is known about protease control in plant-nematode interactions, with only two examples of plant protease inhibition-driven defense being described, although the effectors involved do not contain classical protease inhibitor domains [22,23].

We previously identified 18 novel SCN effector candidates through transcriptome mining using isolated dorsal glands of *H. glycines* and confirmed their expression in the salivary glands by *in situ* hybridization [24]. In the present study, we performed a functional analysis of the *H. schachtii* homolog of one of these candidate effectors, *Hs*2B11, and demonstrate that this effector modulates host immunity. We identified the Arabidopsis Pathogenesis-Related (PR) protein *At*PR-6 (At2g38870), a plant serine protease inhibitor, as an interactor of *Hs*2B11. This interaction was confirmed by split-luciferase assays and bimolecular fluorescent complementation. We show that *At*PR-6 acts as a positive regulator of immunity involved in defense against BCN. Moreover, we identify potential host interactors of *At*PR-6, including plant serine proteases. Our results suggest that *At*PR-6 controls basal immunity through the inhibition of plant serine proteases and/or secreted nematode effectors. This study demonstrates, for the first time, that a cyst-nematode effector targets a protease inhibitor to favor parasitism. These results strengthen our knowledge of plant defense through protease inhibition and suggest that nematodes could counter these defenses through titration of serine protease inhibitors.

## Results

### *Hs*2B11 is a novel effector carrying a unique carboxyl-terminal domain

Bioinformatic analysis of *Hs*2B11 (Hsc_gene_18368) indicates that the gene encodes a putative *H. schachtii* secreted effector with high similarity to GLAND7 (*Hg*2B11; GenBank ID: KJ825718), sharing 85.3% identity at the amino acid level (Fig 1A). We previously localized the expression of GLAND7 transcripts to the dorsal gland of parasitic juveniles using *in situ* hybridization [24]. *Hs*2B11 is also highly expressed in the dorsal gland in transcriptomic analyses conducted on isolated salivary glands (Fig 1B; [25]), and transcripts have been identified during early infection in life cycle RNA-seq analysis data (Fig 1C; [26]). These results suggest that *Hs*2B11 is a secreted effector important for early infection. A BLAST search against WormBase Parasite indicates that this effector is only present in the *H. schachtii* and *H. glycines* genomes and not in closely-related cyst nematodes from the *Globodera* genus, suggesting that this gene has evolved recently. Notably, a cluster of genes encoding paralogues of *Hs*2B11 is present on chromosome 2, suggesting their importance in the infection process (Fig 1D; Fig S1).

**Fig 1.**
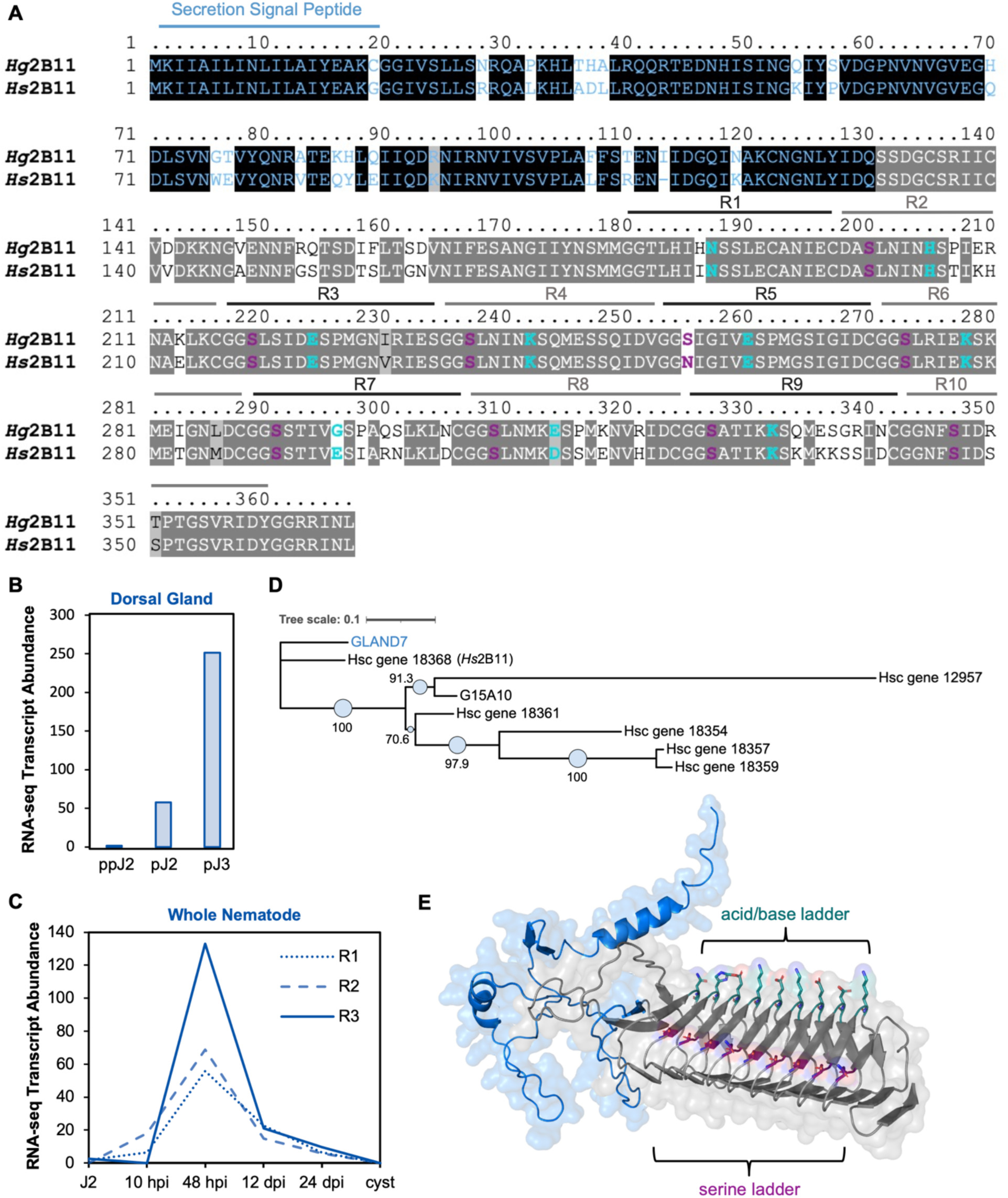
*Hs*2B11 is a predicted effector protein expressed during early infection. (A) Amino acid sequence alignment of 2B11 effectors from *Heterodera glycines* and *Heterodera schachtii* (*Hg*2B11 and *Hs*2B11). The N-terminal domain is indicated in blue text with conserved amino acids in black boxes and the C-terminal domain is identified by black text with conserved residues in gray boxes. The secretion signal peptide (SP) is indicated by a blue line and alternance of repeats (R1-10) in the C-terminal domain are indicated by grey lines. Residues forming acid/base and serine ladders are indicated in turquoise and purple, respectively. (B) RNA-seq expression data from [26] showing the dorsal gland expression of *Hs*2B11 in pre-parasitic J2 (ppJ2), parasitic J2 (pJ2) and parasitic J3 (pJ3) nematodes. (C) RNA-seq expression data from [25] showing the expression of *Hs*2B11 (Hsc_gene_18368) during the life cycle of whole *H. schachtii* nematodes. (D) Phylogenetic tree of the *H. schachtii* and *H. glycines* 2B11 family. The tree scale corresponds to the number of substitutions per site based on the amino-acid matrix (JTT). The genes starting with Hsc_ correspond to *H. schachtii* genes whereas GLAND7 and G15A10 belong to *H. glycines*. (E) 3D structure of *Hs*2B11 as predicted by Alphafold2 (https://github.com/sokrypton/ColabDesign). The N-terminus is indicated in blue and the C-terminus in grey. The alignment of serines or the alternance of acid and basic amino acids are displayed in purple and turquoise, respectively.

The *Hs*2B11 gene was cloned from cDNA collected from parasitic juveniles, corresponding to Hsc_18368 [25]. *Hs*2B11 is a 366 amino acid protein that possesses a predicted secretion signal peptide (according to SignalP analysis; Fig 1A) and lacks a transmembrane domain (according to Phobius analysis). Both *Hs*2B11 and *Hg*2B11 contain two distinct domains, an N-terminal domain with no notable features, and a C-terminal domain that consists of a number of alternating repetitive sequences (Fig 1A; Fig 1E). To explore the significance of this repetitive region, we used Alphafold2 based on colab design [27] to predict its 3D structure (Fig 1E). Even though this software is better suited to predicting already described domains, it predicts a beta-solenoid-like structure stabilized by a hydrophobic core with high confidence (pLDDT score >90). Beta solenoids are a protein fold composed of repeating beta strands, arranged in anti-parallel fashion to form a superhelix. Each repetitive sequence of the C-terminal domain forms one loop, repeating ten times. Interestingly, this structure harbors a serine ladder on one edge, and a ladder of alternating acidic and basic amino acids (asparagine, aspartate, arginine and lysine residues) on another edge, exposing clear polar interaction surfaces (Fig 1E). We ran this structure through the FoldSeek database [28] and obtained significant hits for beta-solenoids, however, these structures varied in the length of repetitive sequences and did not possess the same laddering effect of outward-facing residues on their surfaces, emphasizing the unique structure of this *Hs*2B11 effector.

### *Hs*2B11 localized to the cytoplasm and the endoplasmic reticulum when transiently expressed in *Nicotiana benthamiana*

To gain insight into cellular function of *Hs*2B11, we first studied its subcellular localization *in planta*. We used transient expression in *N. benthamiana* leaves to express GFP fusion constructs with *Hs*2B11 without its endogenous signal peptide and monitored localization using confocal microscopy (Fig 2A). Our observations revealed that the GFP-*Hs*2B11 fusion exhibited a nucleo-cytoplasmic distribution, whereas the *Hs*2B11-eGFP fusion was primarily localized in the endoplasmic reticulum (ER) and the cytoplasm. We validated the ER localization pattern by co-expressing *Hs*2B11-GFP with the ER subcellular marker AtWAK2-mCherry-HDEL (Fig 2B).

**Fig 2.**
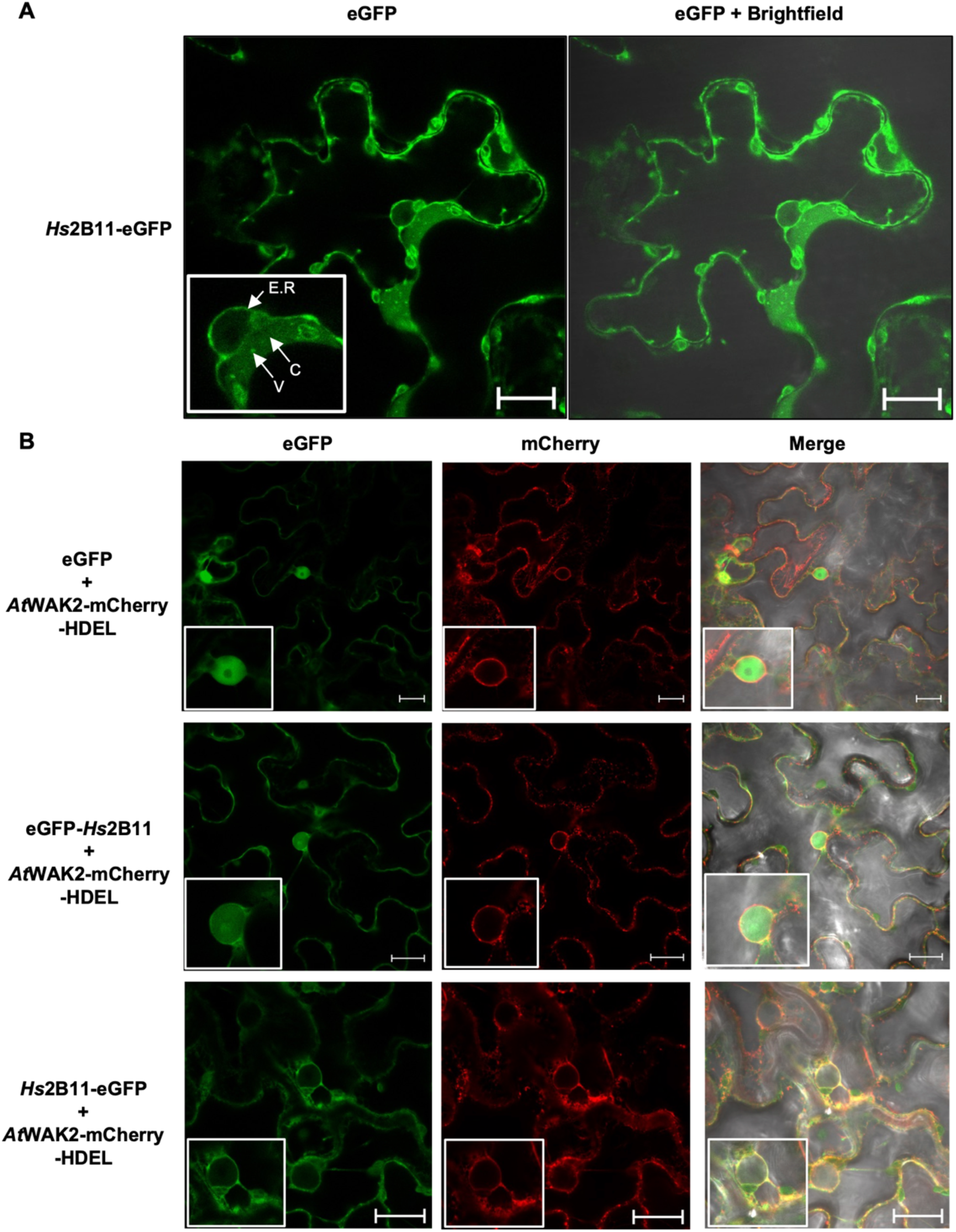
*Hs*2B11 localizes to the cytoplasm and endoplasmic reticulum in *Nicotiana benthamiana.* (A) Subcellular localisation of transiently expressed *Hs*2B11-eGFP in *N. benthamiana* leaves 2 dpi. (B) Co-localization of eGFP-*Hs*2B11 and *Hs*2B11-eGFP with the *A. thaliana* endoplasmic reticulum marker *At*WAK2-mCherry-HDEL. E.R= endoplasmic reticulum; C= cytoplasm; V= vesicle. Experiments were conducted two times with similar results. Bar = 20 µM.

### Expression of *Hs*2B11 in Arabidopsis displays contrasting phenotypes correlated with expression levels

In order to assess if *Hs*2B11 dampens host immunity to promote parasitism we generated two independent Arabidopsis lines (1-3-3 and 14-3-4) expressing the effector under the control of a 35S promoter (Fig 3A). qRT-PCR analysis confirmed the expression of *Hs*2B11 in both transgenic lines (Fig 3B). Line 14-3-4 moderately expressed *Hs*2B11 while line 1-3-3 showed approximately three times higher expression (Fig 3B). Interestingly, moderate expression of *Hs*2B11 resulted in larger rosettes compared to Col-0 plants at 2- and 5-weeks post-germination (wpg) (Fig 3A). In contrast, line 1-3-3, which displayed stronger *Hs2B11* expression, perturbed the growth of rosettes compared to Col-0 and caused spontaneous lesions in leaves (Fig 3A). Consistent with our observations of above-ground growth phenotypes, the highly expressing line (1-3-3) displayed perturbed root growth when grown on MS medium, although no difference in root length was observed in the moderately expressing line (14-3-4) compared to the Col-0, under these conditions (Fig 3C-D). In order to assess the potential role of *Hs*2B11 in immune suppression, we monitored the production of oxidative species in transgenic *Hs*2B11-expressing Arabidopsis lines in response to flg22, an immunogenic peptide derived from bacterial flagellin. We observed that the moderately expressing-*Hs*2B11 line shows a dampened oxidative species burst, whereas the higher expressing-*Hs*2B11 line shows higher oxidative species production (Fig 3E). In contrast, heterologous expression of *Hs*2B11 in Arabidopsis did not impact BCN infection (Fig 3F). As *Hs*2B11 (Fig 1C) and related paralogs (Fig S1) are highly expressed during early infection, we speculate that additional expression in Arabidopsis likely has no observable effect on infection. Together, our results suggest that *Hs*2B11 plays an important role in modulating host defense responses. However, the contrasting immune and physiological phenotypes observed in our two transgenic lines indicates that *Hs*2B11 could interact with plant processes that interplay between defense and growth, in a dose-dependent manner.

**Fig 3.**
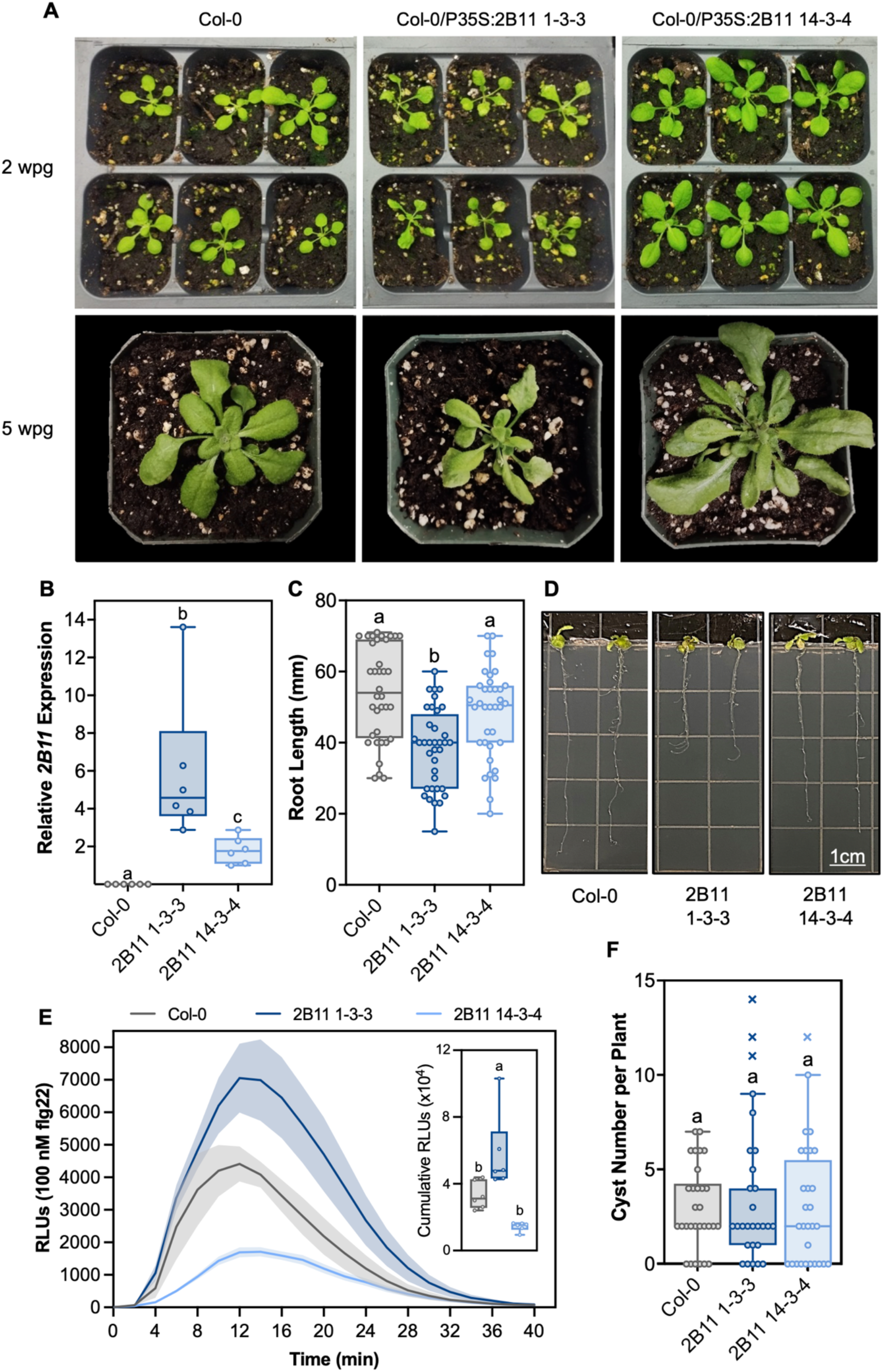
*Hs*2B11 expression in Arabidopsis leads to contrasting growth and immune phenotypes. (A) Photos of two independent transgenic Arabidopsis lines expressing *Hs*2B11 at 2 and 5 weeks post germination (2 wpg and 5 wpg), compared to non-transgenic Columbia (Col-0) controls. (B) Expression levels of *Hs2B11* in transgenic lines assessed by RT-qPCR. Transcripts were normalized to the Arabidopsis housekeeping gene *qUBOX* and compared to Col-0 control plants. (C-D) Root growth of *Hs2B11* expression lines on MS media. (E) Flg22-elicited oxidative species production in *Hs2B11* expressing lines compared to Col-0 controls over a 40 min time course represented as a dynamic curve and cumulative relative light units. (F) Number of cysts per plant in wild-type Col-0 and two *Hs2B11*-expressing lines. Results from two independent experiments are pooled. “x” symbols represent outliers identified using a Grubb’s test. Box plots display interquartile range (25th to 75th percentile) split by a mean line. Whiskers represent maximum and minimum values. Statistically significant values (p<0.05) are indicated by different lower case letters as determined by a one-way analysis of variance followed by Tukey’s post-hoc test. Unless indicated, representative data from one of two experiments is shown.

### *Hs*2B11 interacts with an Arabidopsis PR-6 protein in yeast and *in planta*

In order to gain a better understanding of how *Hs*2B11 is regulating plant defense responses, we sought to identify host interacting proteins by conducting three independent yeast-two hybrid (Y2H) screens against Y2H libraries generated from 3, 7, and 10 dpi BCN-infected Arabidopsis. Screening against the 3-dpi cDNA library yielded 20 potential interacting proteins (Fig 4A; Supplemental File 1). Among these candidate interactors, we identified the Pathogenesis-Related family 6 protein (*At*PR-6; AT2G38870) that was captured 11 times. Using the 7 dpi and 10 dpi libraries, we identified 17 and 26 unique interacting proteins, respectively (Fig 4A; Supplemental File 1). Notably, *At*PR-6 was the only protein to be captured in all three of these screens (Fig 4A). *At*PR-6 belongs to a conserved family of six proteins in Arabidopsis [29], three of which (AT2G38870, AT5G43570 and AT5G43580) are upregulated during BCN infection at 10 dpi (Fig S2). Nevertheless, only one gene (AT2G38870) was identified in our screen. AlphaFold2 predicts that all *At*PR-6 proteins possess a highly conserved C-terminal protease inhibitor domain (Fig S2). In contrast, the N-terminal extension domains were distinct across the six members, which could provide specificity for certain proteases and interactors such as *Hs*2B11.

**Fig 4.**
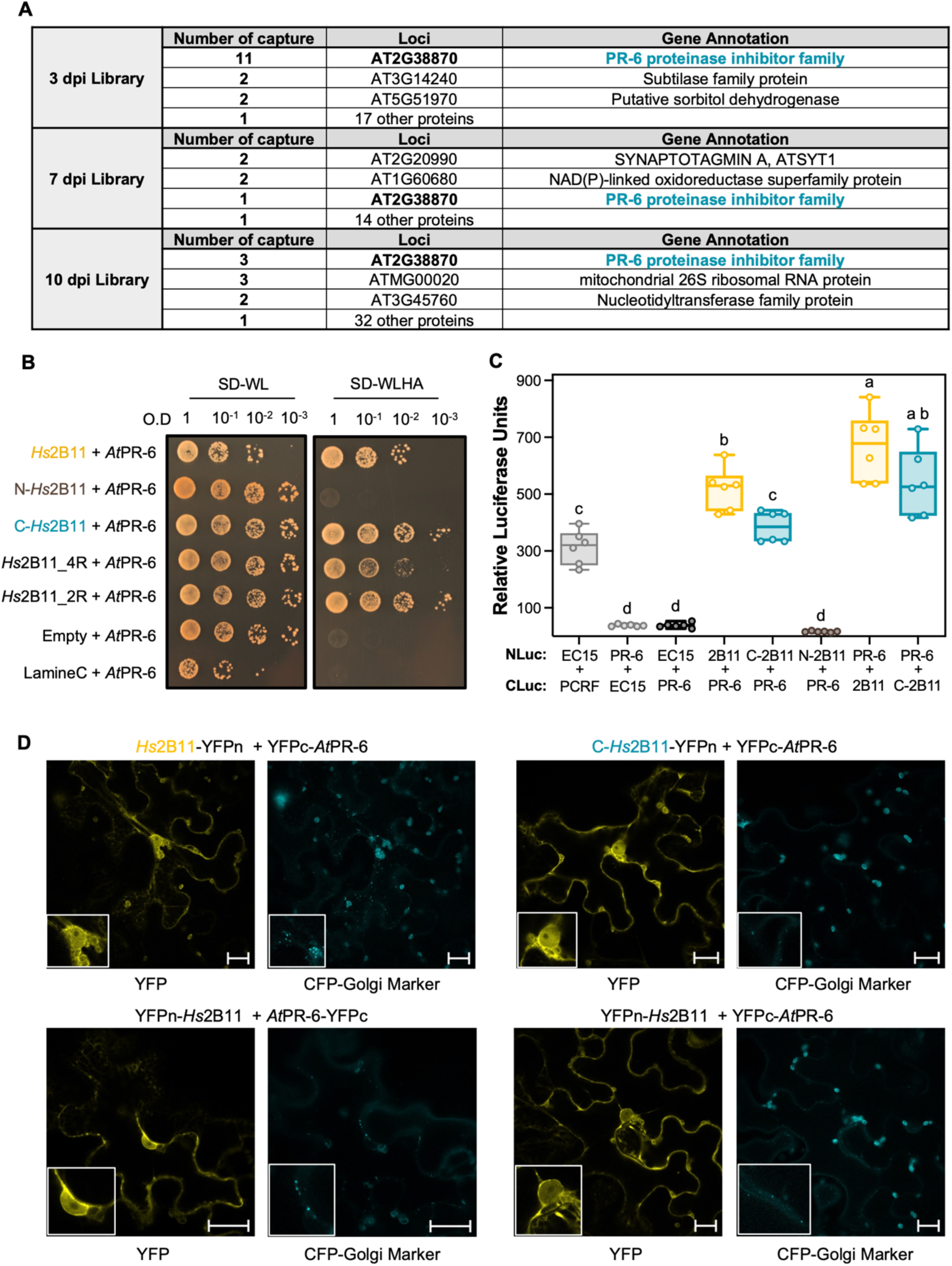
*Hs*2B11 interacts with *At*PR-6 in yeast and *in planta*. (A) Summary of the three independent Y2H screen results against the three libraries of Arabidopsis infected with BCN at 3, 7 and 10 dpi, using *Hs*2B11 as a bait. The number of captures corresponds to the same gene found in independent yeast colonies. (B) Pair-wise Y2H validation of the protein-protein interaction between *Hs*2B11 and *At*PR-6. The N- and C-terminal domains of *Hs*2B11 correspond to the amino acid sequences identified in Fig 1A. *Hs*2B11_2R and *Hs*2B11_4R are composed of 2 or 4 loops of the C-terminal domain of *Hs*2B11. Yeast carrying both vectors (pGBK-T7 and pGAD-T7) are selected on SD-WL media whereas yeast carrying a protein-protein interaction are selected on SD-WLHA, LamineC is used as a negative control. (C) Split-luciferase complementation assay in *N. benthamiana* leaves. The *Phakospora pachyrhizi* effector *Pp*EC15 and its interacting host target, PCRF [60], were used as positive control and *Pp*EC15 with *At*PR-6 was used as a negative control. Lower case letters represent statistically significant groups (p<0.05) determined using a one-way analysis of variance followed by a post-hoc Tukey test. Box plots display interquartile range (25th to 75th percentile) split by a mean line. Whiskers represent maximum and minimum values. Experiments were conducted twice with similar results. (D) BiFC analysis of *Hs*2B11 and *At*PR-6 indicate that the proteins interact in the cytoplasm and endoplasmic reticulum of *N. benthamiana* epidermal cells. Yellow fluorescent protein (YFP) confocal images of BiFC experiments with different combinations of YFPc or YFPn fused, at the C- or N-terminus, to *Hs*2B11 and *At*PR-6, expressed in *N. benthamiana* epidermal cells is shown on the left; and Cyan fluorescent protein (CFP) fused to a Golgi signal peptide on the right. The CFP signal is used to confirm expression of the T-DNA. Bars= 20 µm. BiFC experiments were conducted two times with similar results.

The *At*PR-6 protein family is upregulated in response to biotic stresses, including infection with fungal pathogens [30], exposure to SCN [31], and elicitation with flg22 or the damage-associated molecular pattern (DAMP) *At*Pep1 in roots [32]. *At*PR-6 (AT2G38870) expression has been associated with resistance to *B. cinerea [*30] and *F. graminearum [*33], however, *At*PR-6 has no demonstrated role in nematode defense. Plant extracts containing diverse protease inhibitors were previously shown to inhibit trypsin or subtilases (serine proteases) from various phytopathogen origins such as *Alternaria alternata*, *F. oxysporum* and *B. cinerea* [34]. We therefore aimed to further characterize the role of *At*PR-6 in pathogen defense and its interaction with the effector *Hs*2B11.

We first validated the interaction between *Hs*2B11 and *At*PR-6 using pairwise Y2H analysis (Fig 4B). To determine more specifically the site of interaction between these proteins, we generated N- and C-terminal truncation mutants of *Hs*2B11, corresponding to the domains outlined in Fig 1A. We observed that only the C-terminal repetitive domain of *Hs*2B11 is required for interaction with *At*PR-6 and that the N-terminal domain alone is not capable of interacting with this protein. We further confirmed that the C-terminal domain of *Hs*2B11 is uniquely required for association with *At*PR-6 by performing split-luciferase complementation assays in *N. benthamiana* leaves. In agreement with our Y2H analysis, *Hs*2B11 and *At*PR-6 strongly interact *in planta* (Fig 4C). Notably, there was reduced interaction with the C-terminal domain, compared to the full-length *Hs*2B11 protein, which could indicate that the N-terminal domain partially facilitates interaction between these proteins. Given the repetitive nature of this effector, we were curious if only a subset of these C-terminal repeats is required for interaction. We therefore additionally generated *Hs*2B11 truncation mutants consisting of 2 or 4 repeats, 2R_*Hs*2B11 and 4R_*Hs*2B11, respectively. Indeed, *At*PR-6 associated with both 2R_*Hs*2B11 and 4R_*Hs*2B11 (Fig 4B). Together, these results indicate that *Hs*2B11 strongly associates with *At*PR-6 through interaction with its C-terminal domain.

### *Hs*2B11 and *At*PR-6 interact in the nucleus, cytoplasm and endoplasmic reticulum

To gain insight into the biological relevance of this interaction we conducted bimolecular fluorescence complementation (BiFC) assays to determine the site of interaction of these proteins *in planta*. We identified a strong signal in the ER and cytoplasm of *N. benthamiana* leaf cells transiently expressing *Hs*2B11 and *At*PR-6 possessing N- or C-terminal split-YFP fusions (Fig 4D; Fig S3) which co-localized with the ER marker, AtWAK2 (Fig S4). We performed these assays using a BiFC vector that co-expresses a CFP-tagged Golgi marker in order to confirm the expression of our genes in the absence of YFP re-constitution (Fig 4D). Expression of RFP-*At*PR-6 or *At*PR-6-RFP fusions in *N. benthamiana* resulted in a nucleo-cytoplasmic and nucleo-cytoplasmic/ER localization pattern, respectively (Fig 5). To investigate whether *Hs*2B11 and *At*PR-6 co-localized in the same subcellular compartment, we co-expressed GFP fusions of *Hs*2B11 along with RFP fusions o*f At*PR-6 and observed that they indeed co-localized in the nucleo-cytosolic compartment and/or the ER, depending on the orientation of the tags used (Fig 5).

**Fig 5.**
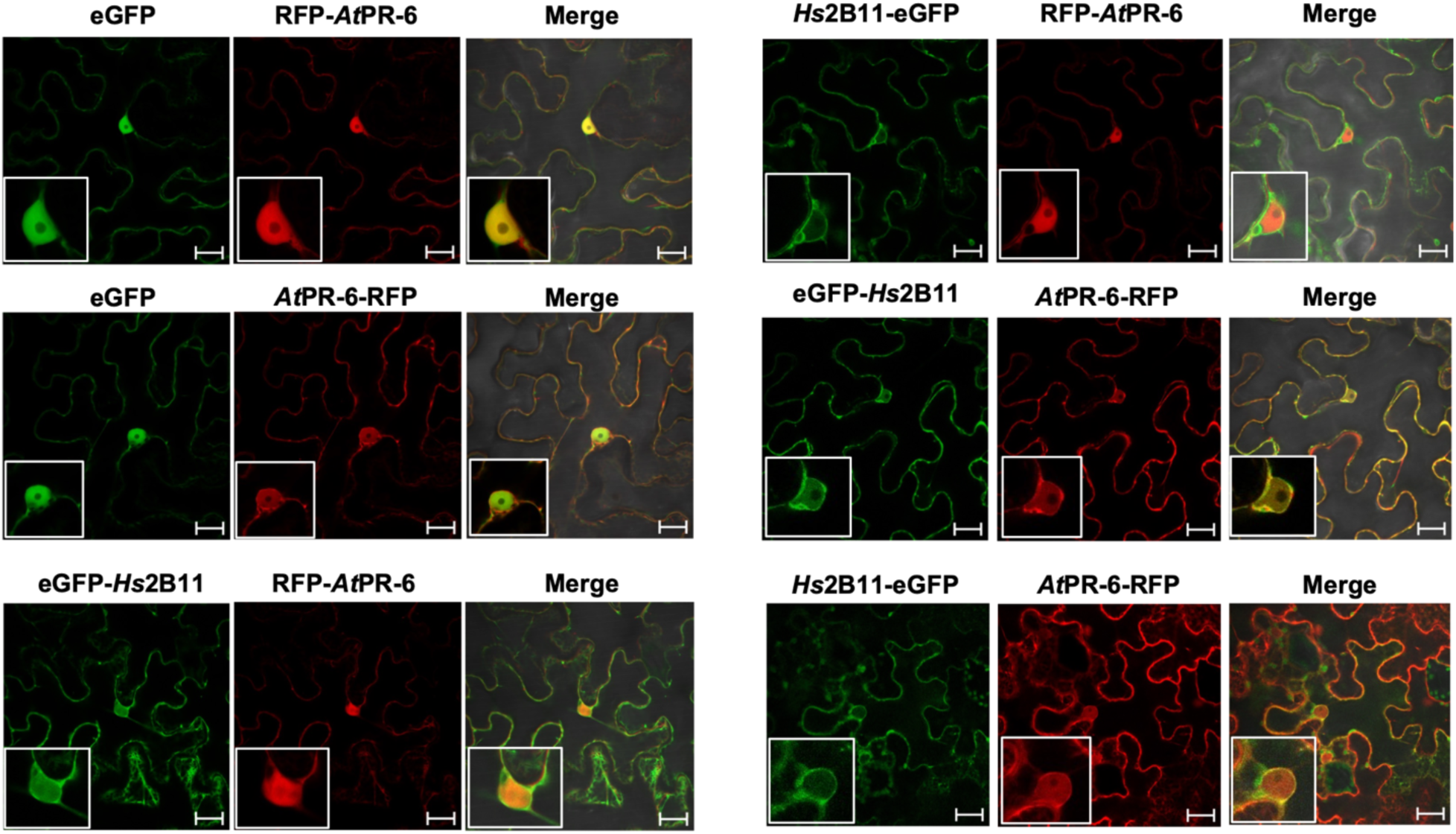
*Hs*2B11 and *At*PR-6 co-localize in the cytoplasm and endoplasmic reticulum in *N. benthamiana.* Co-localization of *Hs*2B11 and *At*PR-6, fused to eGFP and RFP, respectively. eGFP alone is used as a control. Bar = 20 µM. Subcellular localization experiments were conducted two times with similar results.

### *At*PR-6 is upregulated in early syncytia

To investigate the spatio-temporal expression pattern of the *AtPR-6* gene in response to BCN infection, we generated a transcriptional fusion of the 846 bp sequence upstream of the AT2G38870 start codon with the GUS reporter gene (p*AtPR6:GUS*) and produced stable Arabidopsis lines. We performed histochemical GUS assays and found that the *AtPR-6* promoter is active ubiquitously in leaves, trichomes, and roots in the absence of BCN infection, with a stronger expression in the root elongation zone and no expression in the root tip (4 h incubation with X-Gluc) (Fig S5). In order to distinguish changes in expression associated with BCN infection, we repeated these assays using a shorter substrate incubation period (30 min in the presence of X-Gluc). We observed a similar expression pattern in these experiments (Fig 6A-D). We next infected these Arabidopsis lines with *H. schachtii* and monitored GUS expression using this shorter incubation time at 4 dpi and 14 dpi (Fig 6E-H). We observed an upregulation of *At*PR*-6* promoter-driven expression in syncytia at 4 dpi (Fig 6E, 6F) but not at 14 dpi (Fig 6G, 6H), indicating that this gene is upregulated as an early response to BCN infection. This observation is in line with previously established RNA-seq data that indicate an upregulation of *AtPR-6* during early infection with an expression peak at 10 hpi with BCN ([25]; Fig S2).

**Fig 6.**
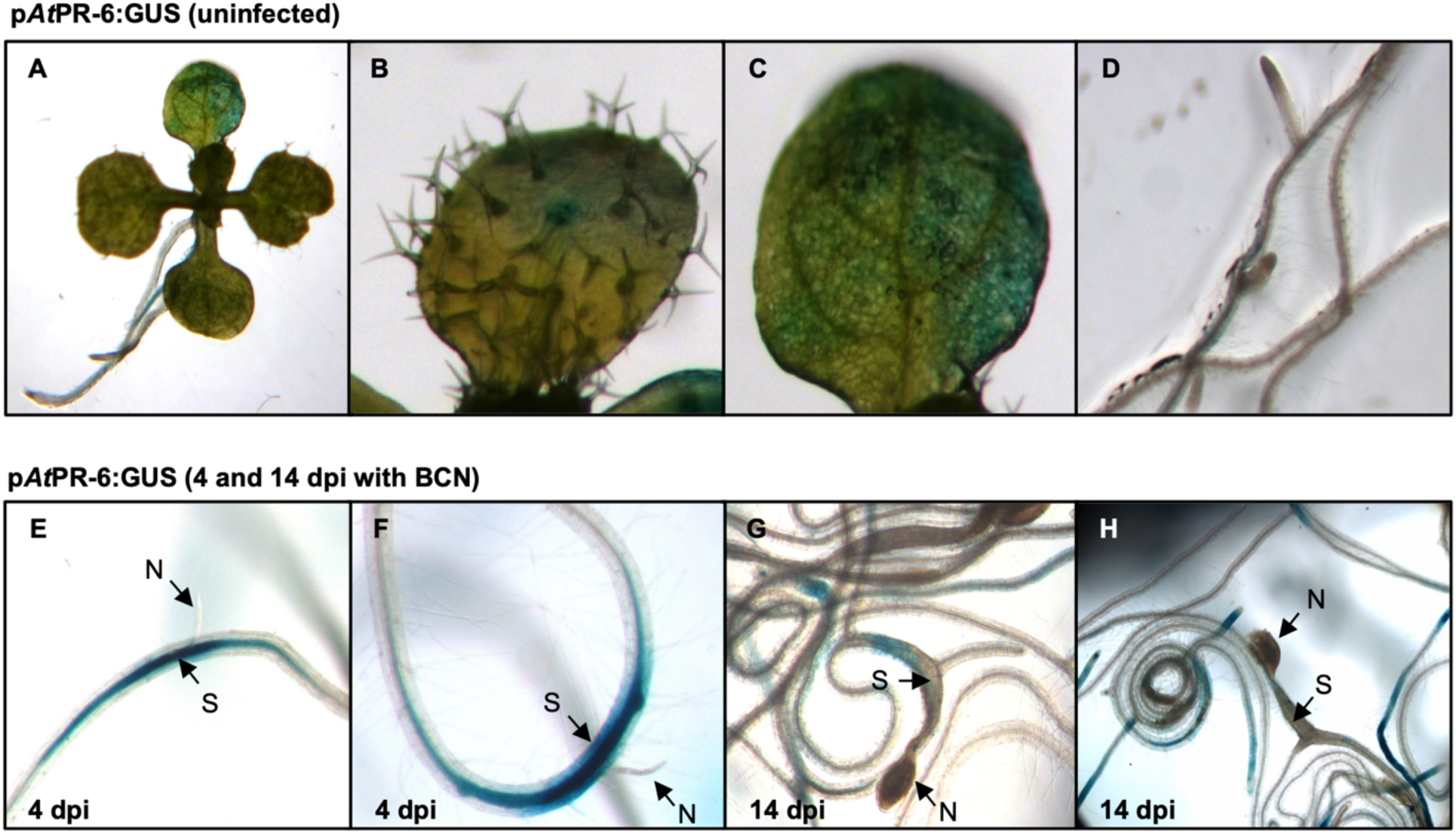
Histochemical localization of GUS activity driven by the *AtPR-6* promoter. GUS activity was observed in (A) uninfected whole Arabidopsis seedlings, (B) emerging leaves with trichomes, (C) leaves and (D) roots. *AtPR-6* promoter activity in BCN-infected Arabidopsis at 4 dpi (E-F) and 14 dpi (G,H). N= nematode; S = syncytium.

### *At*PR-6 is a positive regulator of early immune signaling and provides resistance to *H. schachtii*

Given that our and previous data indicate that *AtPR-6* is responsive to infection and elicitation with immunogenic peptides, it is reasonable to expect the gene has a role in regulating host defense responses. We therefore obtained a T-DNA knockout line of *AtPR-6* (SALK_111051C) (Fig 7A) and conducted infection assays with *H. schachtii*. The *AtPR-6* knockout line was more susceptible to BCN infection, displaying a significantly higher number of cysts per plant compared to wild-type Col-0 plants (Fig 1B). To determine if *AtPR-6* is more generally involved in basal defense, we monitored flg22-induced oxidative species production, which showed a reduced immune response (Fig 1C). Since only one line was available with a T-DNA insertion in the coding region of *At*PR-6, we additionally generated *At*PR-6 overexpression lines driven by a 35S promoter and confirmed *AtPR-6* overexpression by RT-qPCR analysis (Fig 7D). Consistently, *At*PR-6 overexpression lines were more resistant to BCN infection, displaying significantly reduced cysts per plant compared to wild-type Col-0 controls (Fig 1E). Overexpression of *At*PR-6 also correlated with an overproduction of oxidative species in response to flg22 (Fig 1F). Together, our data provide direct evidence for a role of *At*PR-6 in immune signaling and defence against *H. schachtii*.

**Fig 7.**
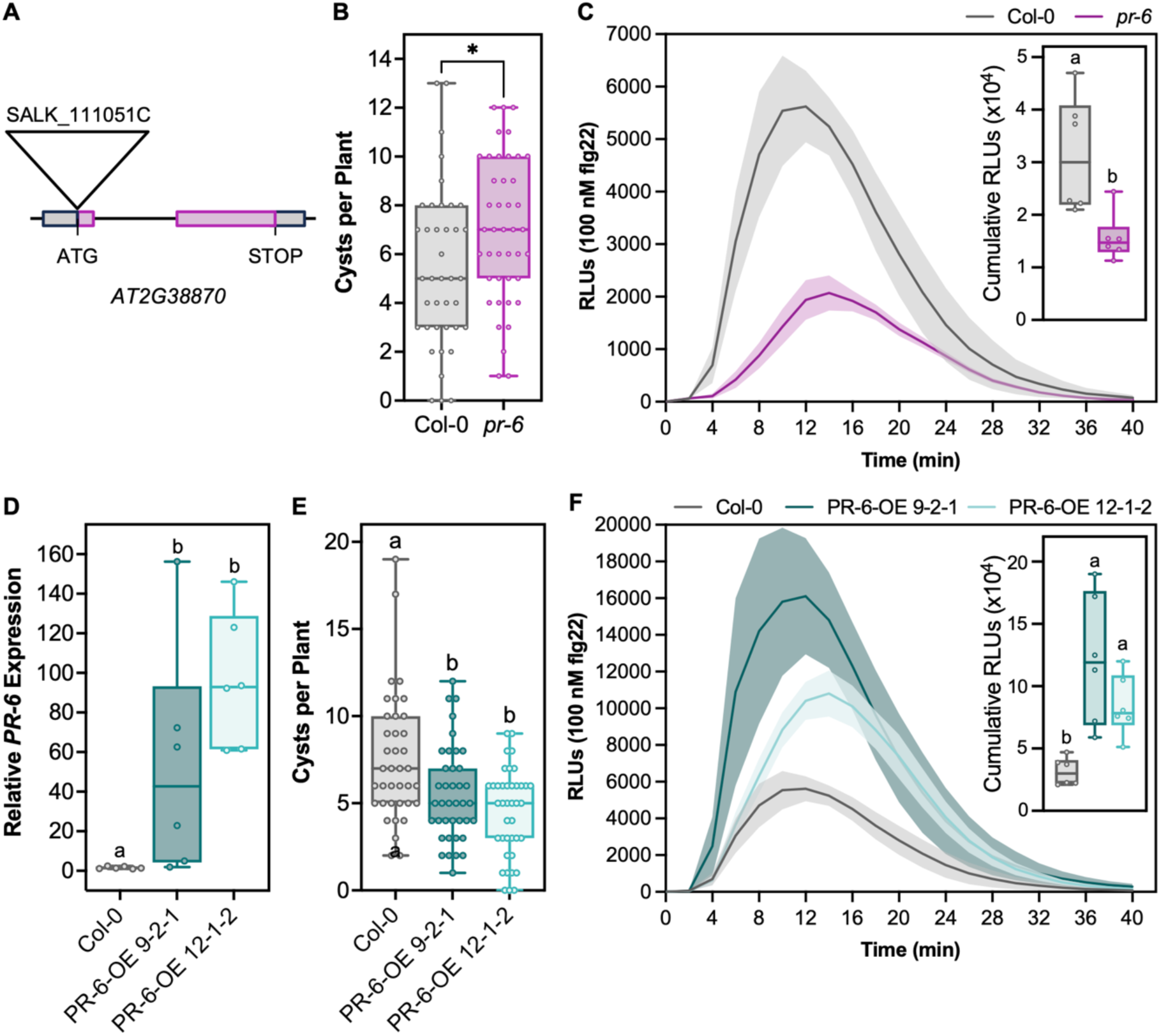
*At*PR-6 is a positive regulator of plant immunity and defense against *H. schachtii* infection. (A) Position of T-DNA insertion of *Atpr-6* knockout allele (Salk_111051C). (B) Number of cyst per plant in Col-0 control line and a *Atpr-6* knockout mutant 4 weeks-post infection with 250 *H. schachtii* second-stage juveniles (J2s). Data from two independent experiments are pooled. Asterisks indicate statistically significant values (p<0.05) as determined by a one-way analysis of variance followed by Tukey’s post-hoc test. (C) Flg22-induced oxidative species production in the Arabidopsis *Atpr-6* mutant line compared to Col-0 control. Representative data from one of two experiments showing similar results are shown. (D) qRT-PCR analysis of two *AtPR-6* Arabidopsis overexpression lines. Experiments were conducted twice with similar results. (E) Number of BCN cysts per plant in Col-0 controls and two *At*PR6 overexpressing lines, 4 weeks-post-infection with 250 *H. schachtii* J2s. (F) ROS burst kinetics in two *At*PR-6 overexpression lines compared to Col-0 controls, after being challenged with flg22. Representative data from one of two experiments with similar results are shown. Box plots display interquartile range (25th to 75th percentile) split by a mean line. Whiskers represent maximum and minimum values. Unless indicated otherwise, statistically significant values (p<0.05) are indicated by different lowercase letters as determined by a one-way analysis of variance followed by Tukey’s post-hoc test.

### *At*PR-6 interacts with plant serine proteases in Y2H

Since little is known about the function of the *At*PR-6 protein encoded by the AT2G38870 gene, we performed a Y2H screen to identify plant interacting proteins. We used *At*PR-6 as bait against a mix of the three cDNA libraries used previously and identified a total of 25 potential targets (Table S1). Among these proteins, we captured two extracellular plant serine proteases (AT1G20160 and AT3G14240), supporting the role of *At*PR-6 as a serine protease inhibitor. Since the cDNA libraries have been prepared from BCN-infected Arabidopsis roots, we also captured three proteins from the nematode, among which were two protein-encoding genes with predicted secretion signal peptides (Hsc_13411 and Hs_G3197) (Table S1). Although these genes are not predicted to encode proteins possessing protease activity, one of these genes (Hsc_20080) is a transthyretin-like protein, which shares similarity to transthyretin proteins that have been demonstrated, in some cases, to possess cryptic serine protease activity [35]. Our data suggest that *At*PR-6 could target both host and nematode serine proteases, in an interplay with growth and defense to counteract nematode infection.

## Discussion

In this study, we functionally characterized *Hs*2B11, a previously uncharacterized BCN effector gene from *H. schachtii*. *Hs*2B11 is one of the multiple pioneer genes that have no described function in the *H. schachtii* genome. Since the *Heterodera* genera have evolved sophisticated relationships with their hosts through the formation of a peculiar feeding site and a narrow host range [36], it is not surprising to find that many pioneer genes have still not been functionally investigated. Multiple paralogues of *Hs*2B11 are found in the genome of *H. schachtii* suggesting that these genes could act redundantly, or have diversified to target a larger range of host targets. Moreover, *Hs*2B11 is an orthologue of GLAND7 and shows sequence and structural similarity to G15A10, an effector also identified in the dorsal gland of SCN [24]. Given that these genes have not been identified outside of the *Heterodera* genera, we speculate that this gene family has evolved recently to act on plant processes related to *Heterodera* cyst nematode infection strategies. Here, we demonstrate that *Hs*2B11 perturbs flg22-induced oxidative species production in Arabidopsis, indicating a role in modulating basal defense responses. We identified an *At*PR-6 protein, belonging to a family of serine protease inhibitors, as an interactor of *Hs*2B11, which positively regulates basal immunity and defense against BCN infection. Our data suggest that *Hs*2B11 targets *At*PR-6 to suppress its endogenous immune functions.

In order to establish a role for *Hs*2B11 in plant parasitism, we generated 35S-driven overexpression lines in Arabidopsis. During the course of experiments, we observed that the level of *Hs2B11* expression in these transgenic lines caused reciprocal phenotypes, with “lower” expression dampening defense and promoting plant growth, and “higher” expression inducing basal defense responses and perturbed plant growth. These results clearly demonstrate a role for *Hs*2B11 in regulating host defense responses that seems to be intricately intertwined with plant growth. This growth-defense trade-off could be the result of targeting a plant protein that interplays between these two processes. This type of expression-dependent reciprocity has been documented previously in the *H. schachtii*-Arabidopsis pathosystem. Low expression of the *H. schachtii* effector 32E03 was shown to inhibit the activity of the histone deacetylase HDT1, causing increased acetylation of histone H3 along the rDNA chromatin, whereas higher expression triggered repression of the same rDNA region [37]. Higher expression also resulted in dramatically smaller plants which were more resistant to BCN infection, whereas lower expression caused no obvious growth defects and higher susceptibility to infection. During a natural infection, effector secretion is tightly regulated to precisely control the quantity and location of specific sets of effectors in the host to optimally promote parasitism. Although ectopic expression of effectors using transgenic systems can provide clues about the function of a potential effector, it does not always reflect its real effect during physiological conditions. We expect that *H. schachtii* would secrete *Hs*2B11 as a means of promoting infection, leading us to hypothesize that the moderately expressing line is the most representative of what would occur during BCN infection. Notably, we did not observe any growth phenotype of *At*PR6 overexpression lines or of the *Atpr6* knockout line, suggesting that *Hs*2B11 could have additional targets with developmental roles. This is also supported by the identification of multiple targets in our Y2H screen.

Nevertheless, *At*PR-6 was identified across the three Y2H libraries suggesting it is a major target of Hs2B11. PR-6 proteins were initially identified by their upregulation during biotic or abiotic stresses [29] and are divided into 17 families based on their function [38]. Members of the *At*PR-6 family are protease inhibitors, with a low molecular mass varying from 8 to 20 kDa, that inhibit proteolytic activities from animals, fungal and vegetal origins [29]. In Arabidopsis, six *AtPR-6* genes have been identified that encode for serine protease inhibitors [38]. The *At*PR-6 protein identified in our screen (At2g38870) encodes for a small (7.6 kDa) potato type I serine protease inhibitor, which has been shown to inhibit the *in vitro* growth of *F. graminearum* through endogenous inhibitor activity [39] and confers resistance to *B. cinerea* when overexpressed in Arabidopsis [30]. One other *At*PR-6 member, At5g43580 (Unusual Serine Protease Inhibitor; UPI), also provides resistance against *B. cinerea* and a generalist herbivore *Trichoplusia ni* [40]. Here, we show that *At*PR-6 positively regulates flg22-mediated oxidative species production, demonstrating a direct role in immune signaling, and provides resistance against BCN infection. Several serine protease inhibitors in this family are also upregulated in soybean in response to SCN infection [41], which could suggest a conserved role in defense against parasitic nematodes.

In order to begin to dissect the role of *At*PR-6 in Arabidopsis we investigated its spatio-temporal localization. GUS reporter-aided promoter analysis indicated that *AtPR-6* is upregulated as an early response to BCN infection, which corresponds to the time frame in which nematodes suppress defense and initiate the formation of the feeding site. Using *pAtPR-6:GUS* reporter lines, we demonstrated that *AtPR-6* gene expression is upregulated in syncytia at early stages of infection (4 dpi), but not at later stages (14 dpi). These results are consistent with RNA-seq expression data of *At*PR-6 during infection [25], with a peak of expression at 10 hpi. These results are also in line with the number of captures of *At*PR-6 in the three different Y2H cDNA libraries using *Hs*2B11 as a bait (11 captures at 3 dpi, 1 capture at 7 dpi and 3 captures at 10 dpi). It is possible that *At*PR-6 could interact with secreted proteases to regulate host defense responses. Indeed, RNA-seq analysis shows a number of effector serine proteases in BCN-infected Arabidopsis during early infection [36]. *Hs*2B11 may have evolved to counteract *At*PR-6-mediated protease inhibition, placing *Hs*2B11 and *At*PR-6 as players in the co-evolution between BCN and their host.

The role of host- and pathogen-encoded serine protease inhibitors in effector-triggered immunity (ETI) have been well-studied [42,43], however their involvement in microbe-associated molecular pattern (MAMP)-triggered responses is not as well known. Since *At*PR-6 has a role in regulating flg22-elicited ROS production in the absence of nematode infection, we expect that it interacts with plant serine proteases in order to execute endogenous functions. Indeed, our Y2H screen uncovered 22 plant candidate *At*PR-6 interactors, among which we found three plant proteases predicted to be found in the apoplast. The first protease is the subtilisin-like serine protease *At*SBT5.2, a serine protease that attenuates the transcriptional activation of defense by the MYB30 transcription factor in Arabidopsis, therefore acting as a negative regulator of immunity [44]. This gene also serves a direct role in attenuating defense by inactivating flagellin immunogenicity through cleavage of the flg22 epitope [45]. Since *At*PR-6 interacts with *At*SBT5.2, and possesses serine protease inhibitor activity, it is feasible that it could inhibit *At*SBT5.2 activity as a means of promoting defense responses. The second plant protease identified as a potential partner of *At*PR-6 is a subtilase family protein, also belonging to a family of serine proteases [46]. We also found a eukaryotic aspartyl protease as a potential interactor of *At*PR-6 which is S-nitrosylated in response to flg22 treatment in Arabidopsis [47], although a direct role in immunity has not been established. We additionally captured three proteins from the nematode that interacted with *At*PR-6 during this screen, two of which are uncharacterized proteins carrying secretion signal peptides, which would be interesting candidates for future investigations. Altogether, our results indicate that *At*PR-6 could interact with endogenous plant proteases in order to promote defense responses in Arabidopsis.

In order to gain structural insight into the interaction between *Hs*2B11 and *At*PR-6, we modeled each of these proteins, individually, using *in silico* folding with AlphaFold2. This is the first instance in which a 3D prediction software has successfully folded an unknown domain in a phytoparasitic nematode effector with such high confidence, emphasizing the need to predict the 3D structure of the entire effectorome in the future to help researchers unravel effector functions. Structural predictions of *Hs*2B11 have provided us clues about how it could interact with *At*PR-6. The intriguing structure of the C-terminal domain of *Hs*2B11, with a tower-like structure that harbors an alignment of serine on one edge, suggests that these serine residues could mimic the serine residue present in the catalytic pocket of serine proteases. This could explain how two to four repetitions alone are able to interact with *At*PR-6. There are two major mechanisms of protease inhibition, irreversible interactions and tight reversible interactions [43]. In irreversible interactions the protease cleaves the protease inhibitor in a way that destroys the catalytic site of protease. As a consequence, the protease inhibitor is sacrificed at the cost of the interactions. In contrast, reversible tight interactions occur by binding to the active site in a substrate-like fashion. This latter type of interaction has been conclusively demonstrated for serine protease inhibitors. We therefore can envision that the C-terminal domain of *Hs*2B11 mimics the active site of a serine protease to temporarily sequester *At*PR-6, titrating its activity as a positive immune regulator in order to favor BCN infection. The involvement of these serine residues in the interaction will need to be further investigated to better understand if and precisely how the effector *Hs*2B11 mechanistically traps *At*PR-6 to dampen plant basal immunity. This type of mechanism has been demonstrated for the *Magnaporthe oryzae* effector AvrPiz-t that targets the trypsin protease inhibitor APIP4 (AvrPiz-t interacting protein 4) and suppresses its activity in order to promote infection [48]. Future work will need to investigate if *Hs*2B11 has a direct effect on *At*PR-6 protease inhibitor activity or if the effector functions by limiting the accumulation of free *At*PR-6 in a sequestering reaction.

In our experiments, we observed that *Hs*2B11 and *At*PR-6 are localized to the cytoplasm and ER, and formed cytoplasmic punctuated structures resembling vesicles, which could indicate that these proteins transit through the plant cell secretion pathway to reach the apoplast. In our BiFC experiments, we also observed that *Hs*2B11 and *At*PR-6 interact in the ER and the cytoplasm. At least two independent studies have shown that the *At*PR-6 encoded by At2g38870 can be identified in the apoplast of Arabidopsis by mass spectrometry [49,50]. Another *At*PR-6 protein, UPI, also accumulates in the extracellular space, specifically in response to methyl jasmonic acid [30]. UPI is also upregulated at 10 hpi with BCN (Fig S2), which could suggest that these proteins act together in early defense against phytoparasitic nematodes. Nevertheless, we did not observe expression of either *Hs*2B11 or *At*PR-6 in the apoplast under our conditions. Future work is needed to determine where *Hs*2B11 localizes during natural infection and whether *At*PR-6 is localized to the apoplast during BCN infection.

Together, our data indicate that *At*PR-6 is a defense protein upregulated in response to BCN infection, possibly to inhibit the action of plant and/or nematode proteases to prevent nematode infection. We hypothesize that *Hs*2B11 is secreted to titrate *At*PR-6 in order to dampen its immune activity in the host during infection. This work provides the first evidence of a cyst-nematode effector targeting a protease inhibitor involved in host defense.

## Materials and methods

### Plasmid Construction

For 35S promoter-driven expression of *Hs*2B11 and *At*PR-6 expression lines, the coding region of *Hs*2B11 (minus signal peptide) or *At*PR-6 was amplified from parasitic BCN cDNA and Arabidopsis cDNA, respectively using gene-specific primers designed to create the BamHI and SstI restriction sites, respectively (Table S2). All PCR amplifications were performed using the Q5 DNA polymerase (New England Biolabs) according to the manufacturer’s instructions. The PCR products were digested, gel purified, ligated into the binary vector pBI121, and verified by Sanger sequencing. For the *At*PR-6 promoter construct, an 846 bp fragment upstream of the start codon of the *At*PR-6 gene was amplified from Arabidopsis genomic DNA using forward and reverse primers containing XbaI and BamHI restriction sites, respectively (Table S2). The purified PCR product was digested by XbaI and BamHI, gel purified, cloned into XbaI-BamHI restriction sites of binary vector pBI101, and confirmed by Sanger sequencing. For subcellular localization or split-luciferase complementation assays, the full-length *Hs*2B11, or its truncation variants, as well as the *At*PR-6 sequences, were amplified using gene specific primers that contain the AttB1 and AttB2 recombination sites. A two-step PCR using AttB adapter primer pairs was used to add the complete AttB recombination sites. The sequences were recombined using the BP Clonase II into a pDONR221 vector and verified by Sanger sequencing. For the subcellular localization, the corresponding recombinant pDONR211 were used in a LR reaction along with the pK7WGF2, pK7FWG2, pK7WGR2, pK7RWG2 destination vectors to produce the *Hs*2B11 and *At*PR-6 fusions with GFP and RFP, respectively, in both orientations. For split-luciferase complementation assays, the corresponding pDONR211 were used in a LR reaction along with the pGWB-nLuc and pGWB-cLuc vectors to produce the *Hs*2B11 and *At*PR-6 fusions with the N- or C-terminal portion of the luciferase gene.

### Generation of Transgenic Arabidopsis Plants

The *Agrobacterium tumefaciens* strain C58 was transformed with the binary plasmids by the freeze-thaw method [51] and used to transform Arabidopsis Col-0 plants using the floral dip method, as described previously by [52]. Transformed T1 plants were screened on Murashige and Skoog (MS) medium containing kanamycin. and homozygous T3 were used for subsequent analysis. The *AtPR-6* knockout line Salk_111051C was obtained from the Salk Institute and homozygous plants were identified by genotyping.

### Plant Materials and Growth Conditions

For ROS burst assays, RT-qPCR analysis and phenotyping, Arabidopsis plants selected on MS plates containing kanamycin were transferred to Sunshine Mix #1-F1P (Sun Gro Horticulture, Agawam, MA, USA) and fertilized with osmocote beads once every two weeks. Plants were grown in a temperature-controlled growth chamber at 22 °C under a 10-h light/14-h dark (150 to 160 μE · m2 · s−1) and no humidity control. For plate-based assays, including BCN infection assays and root length measurements, seeds were surface-disinfected with 70% (v/v) ethanol for 2 minutes, followed by sterilization in a 3% (v/v) sodium hypochlorite solution (Clorox, California, USA) for 15 minutes. Seeds were then rinsed several times with sterile distilled water to remove residual bleach. For the nematode infection assay, sterilized seeds were germinated on modified Knop’s medium solidified with 0.8% (w/v) Daishin agar in 12-well tissue culture plates.

For root measurements, sterilized seeds were sown on square Petri dishes containing half-strength Murashige and Skoog (1/2 MS) medium. To facilitate root growth along the agar surface and allow for clear phenotypic observation, the plates were maintained at a slight incline from the vertical. Root length was measured 21 days after sowing. To ensure sufficient biological replication, 36 plates were utilized for each genotype. *N. benthamiana* plants used for transient expression assays, BiFC and split-luciferase complementation were grown in Sunshine Mix #1-F1P (Sun Gro Horticulture, Agawam, MA, USA), and fertilised with osmocote beads once every two weeks. Plants were under long-day conditions (16-h light/8-h dark) at 24°C.

### GUS Assays

Monitoring of *AtPR6* gene expression was conducted on three-week old, *in vitro* grown Arabidopsis *pAtPR-6:GUS* lines. For expression following BCN infection, plants were inoculated with 250 BCN J2s and collected at 4 and 14 dpi. Histochemical detection of GUS activity was performed according to [53] for 30 min or 4 hours at 37°C in the presence of X-Gluc, as noted in the text.

### Nematode Infection Assays

Ten-day-old seedlings were inoculated with approximately 250 surface-sterilized J2 *H. schachtii* nematodes per plant, as described previously [54]. The inoculated plants were maintained under the same conditions described above for an additional 3 weeks before counting the J4 adult females. Mean values significantly different from that of the wild type were determined in a modified T-test using the statistical software package SAS (P<0.05). Two independent biological replicates were established for each set of conditions.

### RNA extraction and RT-qPCR analysis

Approximately 50 mg of 3-week-old Arabidopsis tissue was used for total RNA isolation using TRIzol reagent (Invitrogen) according to the manufacturer’s instructions. Two µg of RNA and the Maxima First Strand cDNA Synthesis kit with dsDNase (ThermoFisher) were used for cDNA synthesis. A quantitative real-time PCR assay was performed on a 7500 real-time PCR system (Applied Biosystems, CA, USA) using iTaq Universal SYBR Green Supermix (BioRad). For assessment of the overexpression of *Hs2B11* or *AtPR-6*, *AtUBOX* (AT5G15400) was chosen as an internal control to normalize the data. The transcript level was calculated using the 2 ^-ΔΔCT^ method [55]. Primers used for RT-qPCR analysis can be found in Table S2.

### Yeast Two-Hybrid Assays

Y2H screening was performed as described in the BD Matchmaker Library Construction and Screening Kits user manual (Clontech). The coding sequence of *Hs*2B11 o*r At*PR-6 was amplified using forward and reverse primers containing EcoRI and BamHI restriction sites, respectively (Table S2), and fused to the GAL4 DNA-binding domain of pGBKT7 vector to generate pGBKT7-*Hs*2B11 and then introduced into the *Saccharomyces cerevisiae* strain Y187 to generate bait strains. Three Arabidopsis cDNA libraries from roots of ecotype C24 at 3, 7, and 10 dpi after *H. schachtii* infection were generated in *S. cerevisiae* strain AH109 as a fusion to the GAL4 activation domain of pGADT7-Rec2 vector [56]. Screening for interacting proteins and subsequent analyses were performed as described by Clontech protocols with one Y2H screen with each library for *Hs*2B11, and one Y2H screen using a pool of the three libraries for *At*PR-6 bait. To test the potential interaction between full-length or truncated version of *Hs*2B11 and *At*PR-6, the coding sequence of *Hs*2B11 (full-length or truncated) or AtPR-6 was amplified using forward and reverse primers containing EcoRI and BamH1 restriction sites, respectively (Table S2), and fused to the GAL4 DNA activation domain of pGADT7 prey vector to generate pGADT7 recombinant vectors, then introduced into *S. cerevisiae* strain AH109 in combination with the bait vector containing *Hs*2B11 sequences or human Lamin C. The pairwise interaction was tested following Clontech protocols.

### Bimolecular Fluorescence Complementation Assays

The pDOE vectors series have been used to perform the BiFC experiment [57]. The full-length coding sequence of *Hs*2B11 or its C-terminal domain was amplified using specific primers containing NcoI and SpeI. The coding sequence of *At*PR-6 was amplified with specific primers containing KflI and BspEI. The *Hs*2B11 amplicons were digested and cloned into pDOE105, pDOE106, pDOE107 and pDOE108. Subsequently, *At*PR-6 amplicons were digested and cloned into the vectors already containing the *Hs*2B11 full-length or C-terminus domain sequences, or empty vectors to produce the BiFC vectors. Those vectors were transformed into *A. tumefaciens* GV3101 strain through freeze-thaw method. The presence of the insert and vectors were confirmed by colony PCR.

### Transient protein expression and confocal microscopy

Transient expression was achieved by infiltrating *N. benthamiana* leaves with *A. tumefaciens* GV3101 strains harboring GFP- or RFP-fusion or BiFC constructs at a OD (600 nm) between 0.5 and 1, as described previously (REF). Leaves were imaged 48 h after agroinfiltration, with a confocal microscope (LSM700) equipped with an Argon ion and HeNe laser as the excitation source. For simultaneous GFP/RFP imaging, samples were excited at 488 nm for GFP and 551 nm for mRFP, in the multi-track scanning mode. For simultaneous YFP/CFP imaging, samples were excited at 488 nm for GFP and 405 nm for CFP, in the multi-track scanning mode. GFP or YFP emission was detected selectively with a 505–530 nm band-pass emission filter. CFP emission was detected selectively with a 460-500 nm band-pass emission filter. We detected mRFP fluorescence with a 560–615 nm band-pass emission filter. The ER marker pCMU-AtWak2-mCherry-HDEL [58] was used for co-localization studies.

### Split-luciferase complementation assays

Transient expression was achieved by infiltrating *N. benthamiana* leaves with *A. tumefaciens* GV3101 strains harboring split-luciferase constructs at O.D (600 nm) 0.3 each. Three days after infiltration, six discs per construction were collected and incubated with 1 mM luciferin for 15 min in the dark in a 96 well-plate, before reading the luminescence using the SpectraMax spectrophotometer.

### Oxidative Burst Assays

Flg22-elicited ROS production was measured in 4–5-week-old Arabidopsis plants grown on soil and conducted as described previously [59]. Briefly, six leaf discs, per genotype, were collected in a 96-well plate containing 100 µL of water and incubated overnight in the dark at room temperature. The next day water was replaced with a solution containing 100 μM luminol, 10 μg/mL HRP, and 100 nM of flg22 peptide. Readings were taken using a LUM96 module every 2 min over 40 min, with a 1000 ms integration time.

## Author Contributions

JM, PJ, MB and TJB conceived and designed the experiments. JM, MB, AK, PJ, TM and EK performed the experiments. JM, PJ, MB and TJB analyzed the data. JM, MB and TJB wrote the article with input from all authors. TJB and SAW supervised the work. TJB provided funding for the work.

## Acknowledgments

We thank Jackson Goshon for technical assistance.

## Supporting Information Captions

**Fig S1. Expression analysis and structural modeling of *Hs*2B11 paralogs. (**A) RNA-seq expression data from [25] showing the expression of four *Hs*2B11 paralogs during the life cycle of whole *H. schachtii* nematodes. (B) RNA-seq expression data from [26] showing the dorsal gland expression in pre-parasitic J2 (ppJ2), parasitic J2 (pJ2) and parasitic J3 (pJ3) nematodes. (C) 3D structure of *Hs*2B11 paralogs as predicted by AlphafoldII (ColabFold). The N-terminus is indicated in blue and the C-terminus in grey. The alignment of serines or the alternance of acid and basic amino acids are displayed in purple and turquoise, respectively.

**Fig S2**. **Structural modeling and gene expression analysis of *At*PR-6 genes during BCN infection.** (A) AlphaFold II predictions of the six *At*PR-6 genes. (B) RNA-seq expression data from [25] showing the expression of the six *At*PR-6 genes during the life cycle of *H. schachtii* nematodes.

**Fig S3**. **Yellow fluorescent protein (YFP) confocal images of BiFC experiments in *N. benthamiana* epidermal cells.** (A) Empty YFPc vectors and YFPn *Hs*2B11 and (B) the C-terminal fragment of *Hs*2B11, tagged with YFPn and YFPc-tagged *At*PR6. Bars=10 or 20 µm as indicated in the figures. Cyan fluorescent protein (CFP) fused to a Golgi signal peptide on the right. The CFP signal is used to confirm expression of the T-DNA even in the absence of YFP fluorescence.

**Fig S4**. ***Hs*2B11 and *At*PR-6 interact in the ER in *N. benthamiana* epidermal cells**. Co-localization of YFPn-*Hs*2B11 and *At*PR-6-YFPc or YFPc-*At*PR-6 and the ER marker *At*WAK2-mCherrry-HDEL. Bars=10 or 20 µm as indicated in the figures. Cyan fluorescent protein (CFP) fused to a Golgi signal peptide is used to confirm expression of the T-DNA insert.

**Fig S5. Histochemical localization of GUS activity controlled by the *AtPR6* promoter in Arabidopsis after a 4 h incubation with X-Gluc at 37°C.** GUS activity controlled by *AtPR6* promoter in uninfected whole (A) Arabidopsis seedlings, (B) leaves (C) emerging leaves with trichomes, and (D-E) roots.

**Table S1. Y2H summary using *At*PR-6 as a bait.**

**Table S2. Primers used in this study.**

## References

1. Dilawari R, Kaur N, Priyadarshi N, Prakash I, Patra A, Mehta S, et al. Soybean: A key player for global food security. Soybean Improvement. Cham: Springer International Publishing; 2022. pp. 1–46.

2. Rongli, Chen H, Yang Z, Yuan S, Zhou X ’an. Research status of soybean symbiosis nitrogen fixation. Oil Crop Science. 2020;5: 6–10.

3. Phani V, Khan MR, Dutta TK. Plant-parasitic nematodes as a potential threat to protected agriculture: Current status and management options. Crop Prot. 2021;144: 105573.

4. Mitchum MG, Hussey RS, Baum TJ, Wang X, Elling AA, Wubben M, et al. Nematode effector proteins: an emerging paradigm of parasitism. New Phytol. 2013;199: 879–894.

5. Triffitt MJ. Observations on the life-cycle of Heterodera schachtii. J Helminthol. 1930;8: 185–196.

6. Gupta R, Mfarrej MFB, Elnour RO, Hashem M, Ahmad F. Defence response of host plants for cyst nematode: A review on parasitism and defence. J King Saud Univ Sci. 2023;35: 102829.

7. Hewezi T, Baum TJ. Manipulation of plant cells by cyst and root-knot nematode effectors. Mol Plant Microbe Interact. 2013;26: 9–16.

8. Wang Y, Wang Y, Wang Y. Apoplastic Proteases: Powerful Weapons against Pathogen Infection in Plants. Plant Commun. 2020;1: 100085.

9. Figaj D, Ambroziak P, Przepiora T, Skorko-Glonek J. The Role of Proteases in the Virulence of Plant Pathogenic Bacteria. Int J Mol Sci. 2019;20. doi:10.3390/ijms20030672

10. Chandrasekaran M, Thangavelu B, Chun SC, Sathiyabama M. Proteases from phytopathogenic fungi and their importance in phytopathogenicity. J Gen Plant Pathol. 2016;82: 233–239.

11. Vieira P, Danchin EGJ, Neveu C, Crozat C, Jaubert S, Hussey RS, et al. The plant apoplasm is an important recipient compartment for nematode secreted proteins. J Exp Bot. 2011;62: 1241–1253.

12. Zhang J, Li W, Xiang T, Liu Z, Laluk K, Ding X, et al. Receptor-like cytoplasmic kinases integrate signaling from multiple plant immune receptors and are targeted by a Pseudomonas syringae effector. Cell Host Microbe. 2010;7: 290–301.

13. Qi D, Dubiella U, Kim SH, Sloss DI, Dowen RH, Dixon JE, et al. Recognition of the protein kinase AVRPPHB SUSCEPTIBLE1 by the disease resistance protein RESISTANCE TO PSEUDOMONAS SYRINGAE5 is dependent on s-acylation and an exposed loop in AVRPPHB SUSCEPTIBLE1. Plant Physiol. 2014;164: 340–351.

14. Bellafiore S, Shen Z, Rosso M-N, Abad P, Shih P, Briggs SP. Direct identification of the Meloidogyne incognita secretome reveals proteins with host cell reprogramming potential. PLoS Pathog. 2008;4: e1000192.

15. Hu L-J, Wu X-Q, Ding X-L, Ye J-R. Comparative transcriptomic analysis of candidate effectors to explore the infection and survival strategy of Bursaphelenchus xylophilus during different interaction stages with pine trees. BMC Plant Biol. 2021;21: 224.

16. Haegeman A, Mantelin S, Jones JT, Gheysen G. Functional roles of effectors of plant-parasitic nematodes. Gene. 2012;492: 19–31.

17. Margets A, Foster J, Kumar A, Maier TR, Masonbrink R, Mejias J, et al. The Soybean Cyst Nematode Effector Cysteine Protease 1 (CPR1) Targets a Mitochondrial Soybean Branched-Chain Amino Acid Aminotransferase (GmBCAT1). Mol Plant Microbe Interact. 2024;37: 751–764.

18. Pekkarinen AI, Longstaff C, Jones BL. Kinetics of the inhibition of fusarium serine proteinases by barley (Hordeum vulgare L.) inhibitors. J Agric Food Chem. 2007;55: 2736–2742.

19. A Bowman–Birk-Type Trypsin-Chymotrypsin Inhibitor from Broad Beans. Biochem Biophys Res Commun. 2001;289: 91–96.

20. Liu Y, Gong T, Kong X, Sun J, Liu L. XYLEM CYSTEINE PEPTIDASE 1 and its inhibitor CYSTATIN 6 regulate pattern-triggered immunity by modulating the stability of the NADPH oxidase RBOHD. Plant Cell. 2023; koad262.

21. Song J, Win J, Tian M, Schornack S, Kaschani F, Ilyas M, et al. Apoplastic effectors secreted by two unrelated eukaryotic plant pathogens target the tomato defense protease Rcr3. Proc Natl Acad Sci U S A. 2009;106: 1654–1659.

22. Lozano-Torres JL, Wilbers RHP, Warmerdam S, Finkers-Tomczak A, Diaz-Granados A, van Schaik CC, et al. Apoplastic venom allergen-like proteins of cyst nematodes modulate the activation of basal plant innate immunity by cell surface receptors. PLoS Pathog. 2014;10: e1004569.

23. Pogorelko GV, Juvale PS, Rutter WB, Hütten M, Maier TR, Hewezi T, et al. Re-targeting of a plant defense protease by a cyst nematode effector. Plant J. 2019;98: 1000–1014.

24. Noon JB, Hewezi T, Maier TR, Simmons C, Wei J-Z, Wu G, et al. Eighteen New Candidate Effectors of the Phytonematode Heterodera glycines Produced Specifically in the Secretory Esophageal Gland Cells During Parasitism. Phytopathology. 2015;105: 1362–1372.

25. Siddique S, Radakovic ZS, Hiltl C, Pellegrin C, Baum TJ, Beasley H, et al. The genome and lifestage-specific transcriptomes of a plant-parasitic nematode and its host reveal susceptibility genes involved in trans-kingdom synthesis of vitamin B5. Nat Commun. 2022;13: 6190.

26. Molloy B, Shin DS, Long J, Pellegrin C, Senatori B, Vieira P, et al. The origin, deployment, and evolution of a plant-parasitic nematode effectorome. PLoS Pathog. 2024;20: e1012395.

27. Jumper J, Evans R, Pritzel A, Green T, Figurnov M, Ronneberger O, et al. Highly accurate protein structure prediction with AlphaFold. Nature. 2021;596: 583–589.

28. van Kempen M, Kim SS, Tumescheit C, Mirdita M, Lee J, Gilchrist CLM, et al. Fast and accurate protein structure search with Foldseek. Nat Biotechnol. 2023. doi:10.1038/s41587-023-01773-0

29. Sels J, Mathys J, De Coninck BMA, Cammue BPA, De Bolle MFC. Plant pathogenesis-related (PR) proteins: a focus on PR peptides. Plant Physiol Biochem. 2008;46: 941–950.

30. Chassot C, Nawrath C, Métraux J-P. Cuticular defects lead to full immunity to a major plant pathogen. Plant J. 2007;49: 972–980.

31. Alkharouf NW, Klink VP, Chouikha IB, Beard HS, MacDonald MH, Meyer S, et al. Timecourse microarray analyses reveal global changes in gene expression of susceptible Glycine max (soybean) roots during infection by Heterodera glycines (soybean cyst nematode). Planta. 2006;224: 838–852.

32. Igarashi D, Bethke G, Xu Y, Tsuda K, Glazebrook J, Katagiri F. Pattern-triggered immunity suppresses programmed cell death triggered by fumonisin b1. PLoS One. 2013;8: e60769.

33. Chiu T, Poucet T, Li Y. The potential of plant proteins as antifungal agents for agricultural applications. Synth Syst Biotechnol. 2022;7: 1075–1083.

34. Dunaevskiĭ IE, Tsybina TA, Beliakova GA, Domash VI, Shapno TP, Zabreĭko SA, et al. [Proteinase inhibitors as antistress proteins in higher plants]. Prikl Biokhim Mikrobiol. 2005;41: 392–396.

35. Liz MA, Faro CJ, Saraiva MJ, Sousa MM. Transthyretin, a new cryptic protease. J Biol Chem. 2004;279: 21431–21438.

36. Siddique S, Coomer A, Baum T, Williamson VM. Recognition and Response in Plant-Nematode Interactions. Annu Rev Phytopathol. 2022;60: 143–162.

37. Vijayapalani P, Hewezi T, Pontvianne F, Baum TJ. An Effector from the Cyst Nematode Derepresses Host rRNA Genes by Altering Histone Acetylation. Plant Cell. 2018;30: 2795–2812.

38. Ali S, Ganai BA, Kamili AN, Bhat AA, Mir ZA, Bhat JA, et al. Pathogenesis-related proteins and peptides as promising tools for engineering plants with multiple stress tolerance. Microbiol Res. 2018;212–213: 29–37.

39. Bleackley MR, Dawson CS, McKenna JA, Quimbar P, Hayes BME, van der Weerden NL, et al. Synergistic Activity between Two Antifungal Proteins, the Plant Defensin NaD1 and the Bovine Pancreatic Trypsin Inhibitor. mSphere. 2017;2. doi:10.1128/mSphere.00390-17

40. Laluk K, Mengiste T. The Arabidopsis extracellular UNUSUAL SERINE PROTEASE INHIBITOR functions in resistance to necrotrophic fungi and insect herbivory. Plant J. 2011;68: 480–494.

41. Khan R, Alkharouf N, Beard H, Macdonald M, Chouikha I, Meyer S, et al. Microarray analysis of gene expression in soybean roots susceptible to the soybean cyst nematode two days post invasion. J Nematol. 2004;36: 241–248.

42. Jashni MK, Mehrabi R, Collemare J, Mesarich CH, de Wit PJGM. The battle in the apoplast: further insights into the roles of proteases and their inhibitors in plant-pathogen interactions. Front Plant Sci. 2015;6: 584.

43. Clemente M, Corigliano MG, Pariani SA, Sánchez-López EF, Sander VA, Ramos-Duarte VA. Plant Serine Protease Inhibitors: Biotechnology Application in Agriculture and Molecular Farming. Int J Mol Sci. 2019;20. doi:10.3390/ijms20061345

44. Serrano I, Buscaill P, Audran C, Pouzet C, Jauneau A, Rivas S. A non canonical subtilase attenuates the transcriptional activation of defence responses in Arabidopsis thaliana. Elife. 2016;5. doi:10.7554/eLife.19755

45. Buscaill P, Sanguankiattichai N, Kaschani F, Huang J, Mooney BC, Li Y, et al. Subtilase SBT5.2 inactivates flagellin immunogenicity in the plant apoplast. Nat Commun. 2024;15: 10431.

46. Schaller A, Stintzi A, Rivas S, Serrano I, Chichkova NV, Vartapetian AB, et al. From structure to function - a family portrait of plant subtilases. New Phytol. 2018;218: 901–915.

47. Lawrence SR 2nd, Gaitens M, Guan Q, Dufresne C, Chen S. S-Nitroso-Proteome Revealed in Stomatal Guard Cell Response to Flg22. Int J Mol Sci. 2020;21. doi:10.3390/ijms21051688

48. Zhang C, Fang H, Shi X, He F, Wang R, Fan J, et al. A fungal effector and a rice NLR protein have antagonistic effects on a Bowman-Birk trypsin inhibitor. Plant Biotechnol J. 2020;18: 2354–2363.

49. Boudart G, Jamet E, Rossignol M, Lafitte C, Borderies G, Jauneau A, et al. Cell wall proteins in apoplastic fluids of Arabidopsis thaliana rosettes: identification by mass spectrometry and bioinformatics. Proteomics. 2005;5: 212–221.

50. Jiang S, Pan L, Zhou Q, Xu W, He F, Zhang L, et al. Analysis of the apoplast fluid proteome during the induction of systemic acquired resistance in Arabidopsis thaliana. PeerJ. 2023;11: e16324.

51. Höfgen R, Willmitzer L. Storage of competent cells for Agrobacterium transformation. Nucleic Acids Res. 1988;16: 9877.

52. Clough SJ, Bent AF. Floral dip: a simplified method for Agrobacterium-mediated transformation of Arabidopsis thaliana. Plant J. 1998;16: 735–743.

53. Jefferson RA, Kavanagh TA, Bevan MW. GUS fusions: beta-glucuronidase as a sensitive and versatile gene fusion marker in higher plants. EMBO J. 1987;6: 3901–3907.

54. Baum TJ, Wubben MJ, Hardyy KA, Su H, Rodermel SR. A Screen for Arabidopsis thaliana Mutants with Altered Susceptibility to Heterodera schachtii. J Nematol. 2000;32: 166–173.

55. Livak KJ, Schmittgen TD. Analysis of relative gene expression data using real-time quantitative PCR and the 2(-Delta Delta C(T)) Method. Methods. 2001;25: 402–408.

56. Hewezi T, Howe P, Maier TR, Hussey RS, Mitchum MG, Davis EL, et al. Cellulose binding protein from the parasitic nematode Heterodera schachtii interacts with Arabidopsis pectin methylesterase: cooperative cell wall modification during parasitism. Plant Cell. 2008;20: 3080–3093.

57. Gookin TE, Assmann SM. Significant reduction of BiFC non-specific assembly facilitates in planta assessment of heterotrimeric G-protein interactors. Plant J. 2014;80: 553–567.

58. Ivanov S, Harrison MJ. A set of fluorescent protein-based markers expressed from constitutive and arbuscular mycorrhiza-inducible promoters to label organelles, membranes and cytoskeletal elements in Medicago truncatula. Plant J. 2014;80: 1151–1163.

59. Bredow M, Sementchoukova I, Siegel K, Monaghan J. Pattern-Triggered Oxidative Burst and Seedling Growth Inhibition Assays in Arabidopsis thaliana. JoVE (Journal of Visualized Experiments). 2019; e59437.

60. Chicowski AS, Qi M, Variz H, Bredow M, Montes-Serey C, Caiazza F, et al. A soybean rust effector protease suppresses host immunity and cleaves a 3-deoxy-7-phosphoheptulonate synthase. bioRxiv. 2023. doi:10.1101/2023.09.07.556260

