## Supplemental Information for "The sugar-beet cyst nematode effector *Hs*2B11 targets the Arabidopsis serine protease inhibitor *At*PR-6 to favor parasitism"

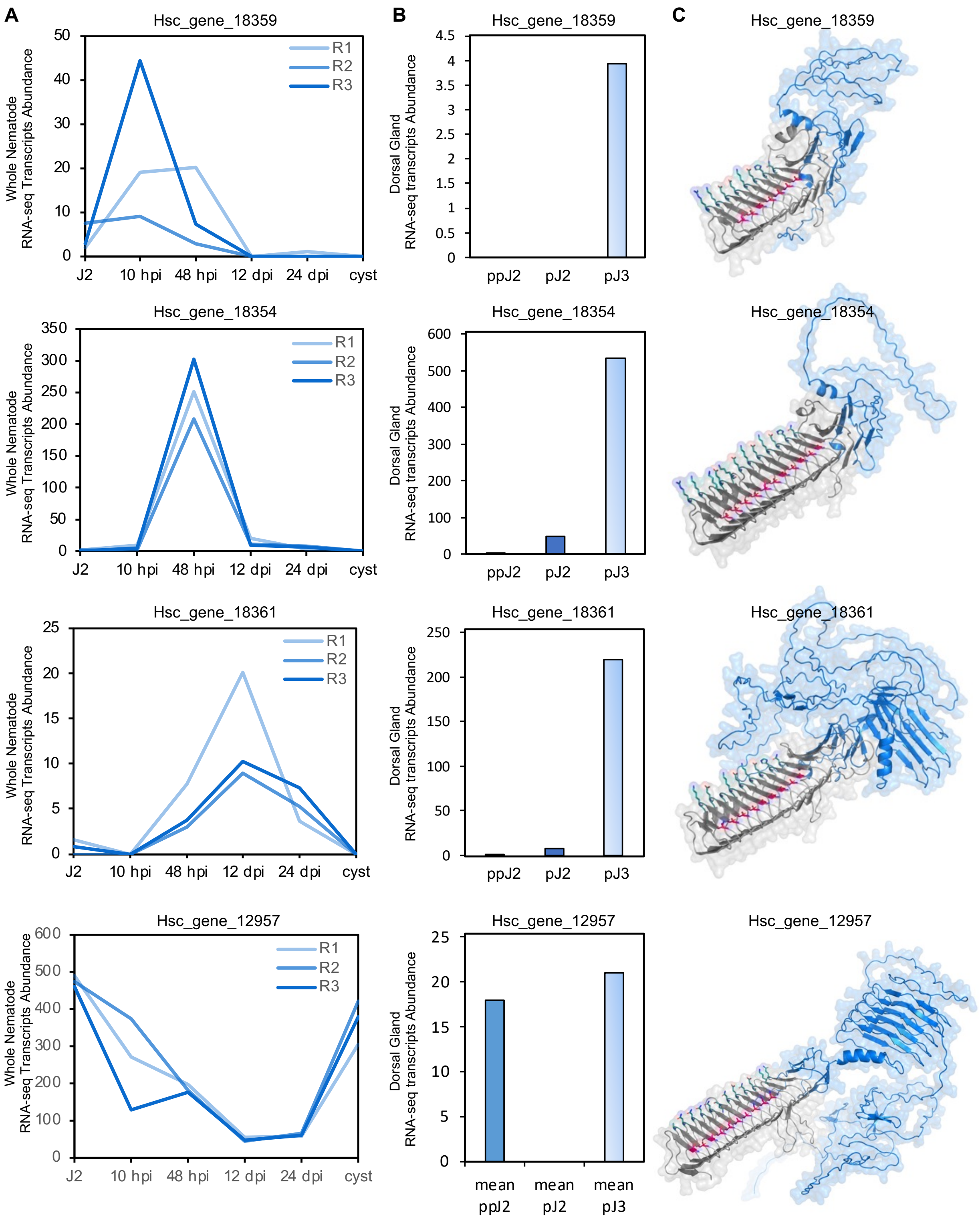
A
B
C
Hsc_gene_18359
Hsc_gene_18354
Hsc_gene_18361
Hsc_gene_12957
Whole Nematode RNA-seq Transcripts Abundance
Dorsal Gland RNA-seq transcripts Abundance
R1
R2
R3
J2
10 hpi
48 hpi
12 dpi
24 dpi
cyst
ppJ2
pJ2
pJ3
mean ppJ2
mean pJ2
mean pJ3

**Fig S1. Expression analysis and structural modeling of *Hs*2B11 paralogs. (**A) RNA-seq expression data from [25] showing the expression of four *Hs*2B11 paralogs during the life cycle of whole *H. schachtii* nematodes*.* (B) RNA-seq expression data from [26] showing the dorsal gland expression in pre-parasitic J2 (ppJ2), parasitic J2 (pJ2) and parasitic J3 (pJ3) nematodes*.* (C) 3D structure of *Hs*2B11 paralogs as predicted by AlphafoldII (ColabFold). The N-terminus is indicated in blue and the C-terminus in grey. The alignment of serines or the alternance of acid and basic amino acids are displayed in purple and turquoise, respectively.

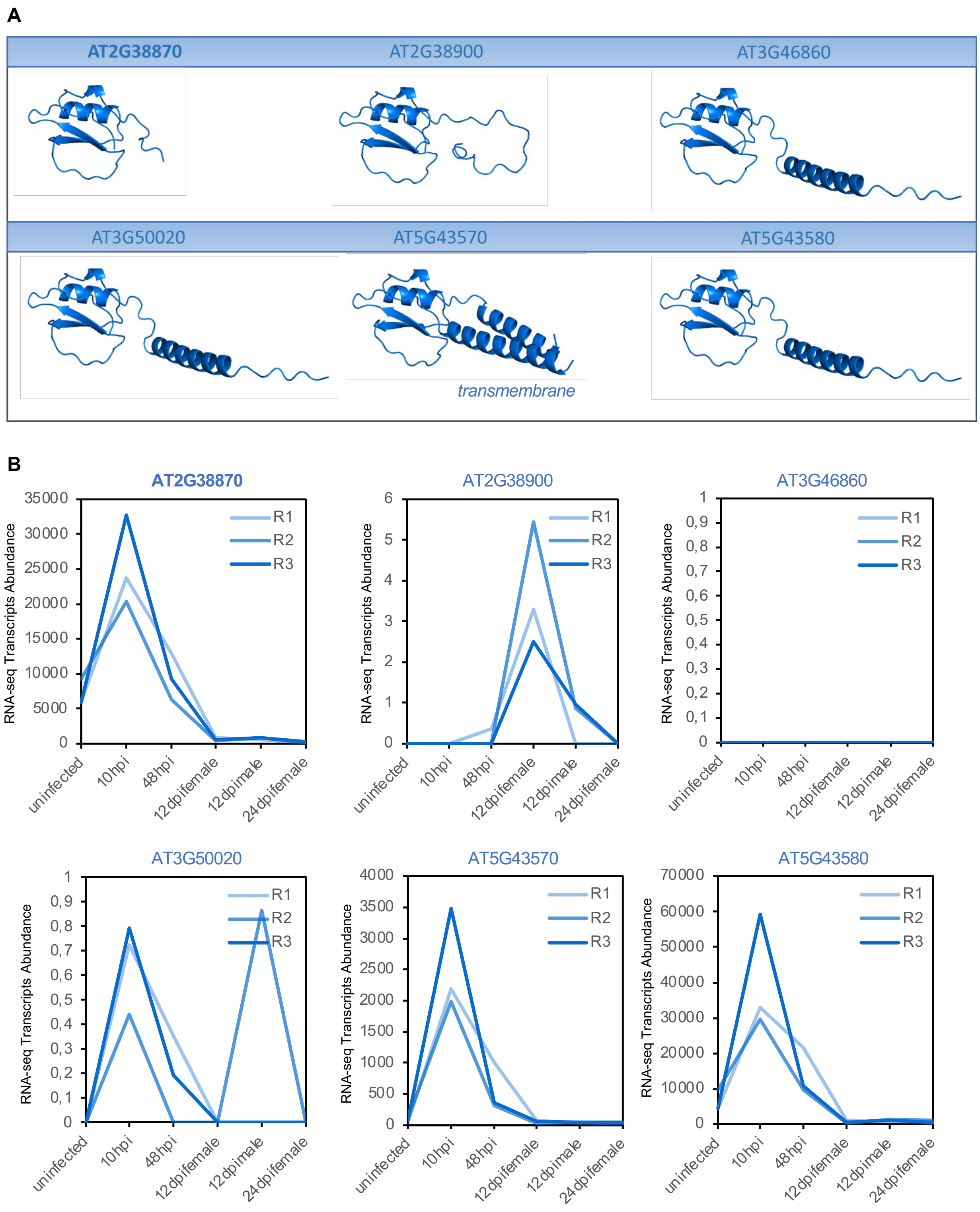


**Fig S2**. **Structural modeling and gene expression analysis of *At*PR-6 genes during BCN infection.** (A) AlphaFold II predictions of the six *At*PR-6 genes. (B) RNA-seq expression data from [25] showing the expression of the six *At*PR-6 genes during the life cycle of *H. schachtii* nematodes.

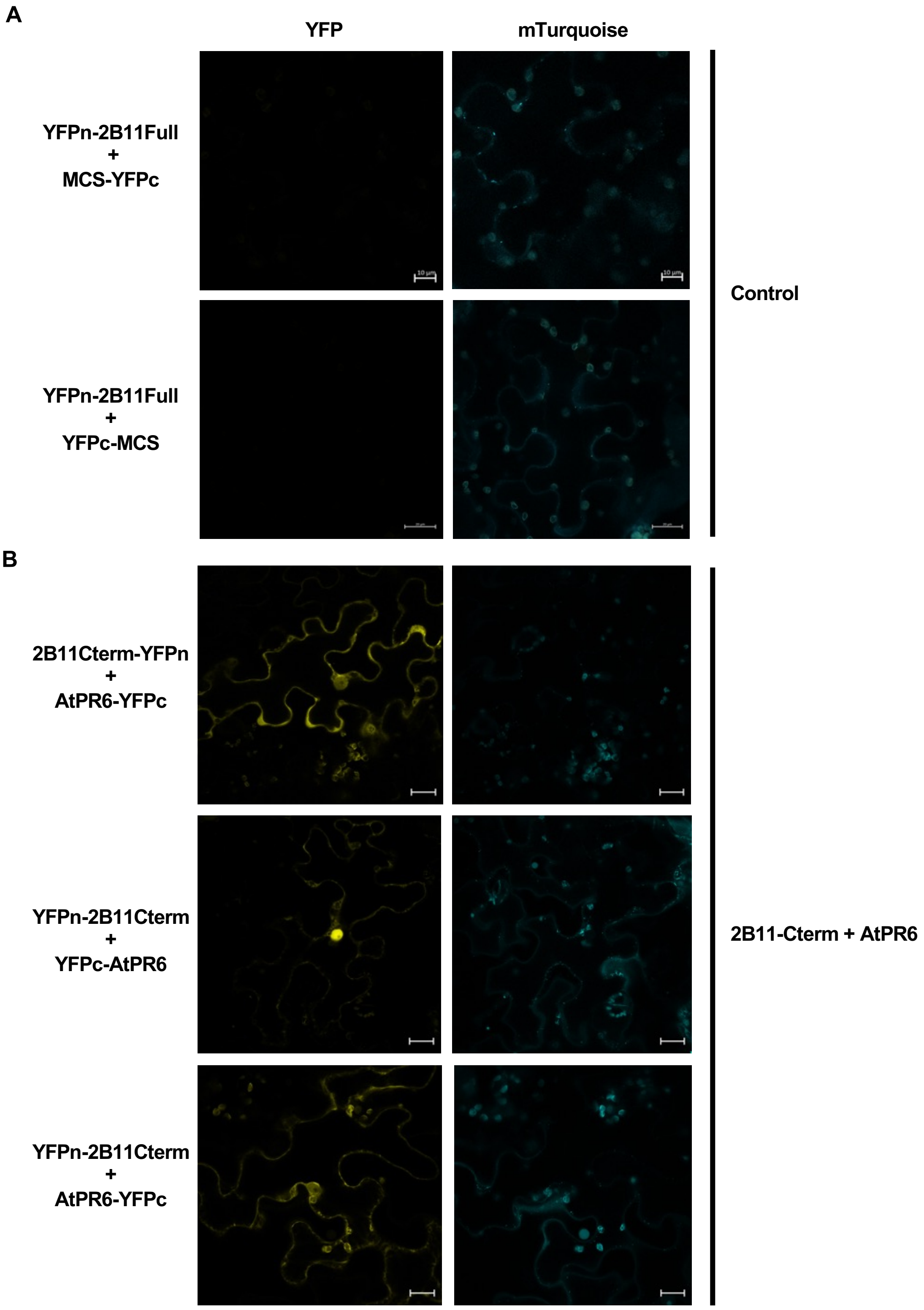
A
YFP
mTurquoise
YFPn-2B11Full
+
MCS-YFPc
YFPn-2B11Full
+
YFPc-MCS
Control
B
2B11Cterm-YFPn
+
AtPR6-YFPc
YFPn-2B11Cterm
+
YFPc-AtPR6
YFPn-2B11Cterm
+
AtPR6-YFPc
2B11-Cterm + AtPR6

**Fig S3**. Yellow fluorescent protein (YFP) confocal images of BiFC experiments expressed in *N. benthamiana* epidermal cells using (A) empty YFPc vectors and YFPn *Hs*2B11 and (B) the C-terminal fragment of *Hs*2B11, tagged with YFPn and YFPc-tagged *At*PR6. Bars=10 or 20 μm as indicated in the figures. Cyan fluorescent protein (CFP) fused to a Golgi signal peptide on the right. The CFP signal is used to confirm expression of the T-DNA even in the absence of YFP fluorescence.

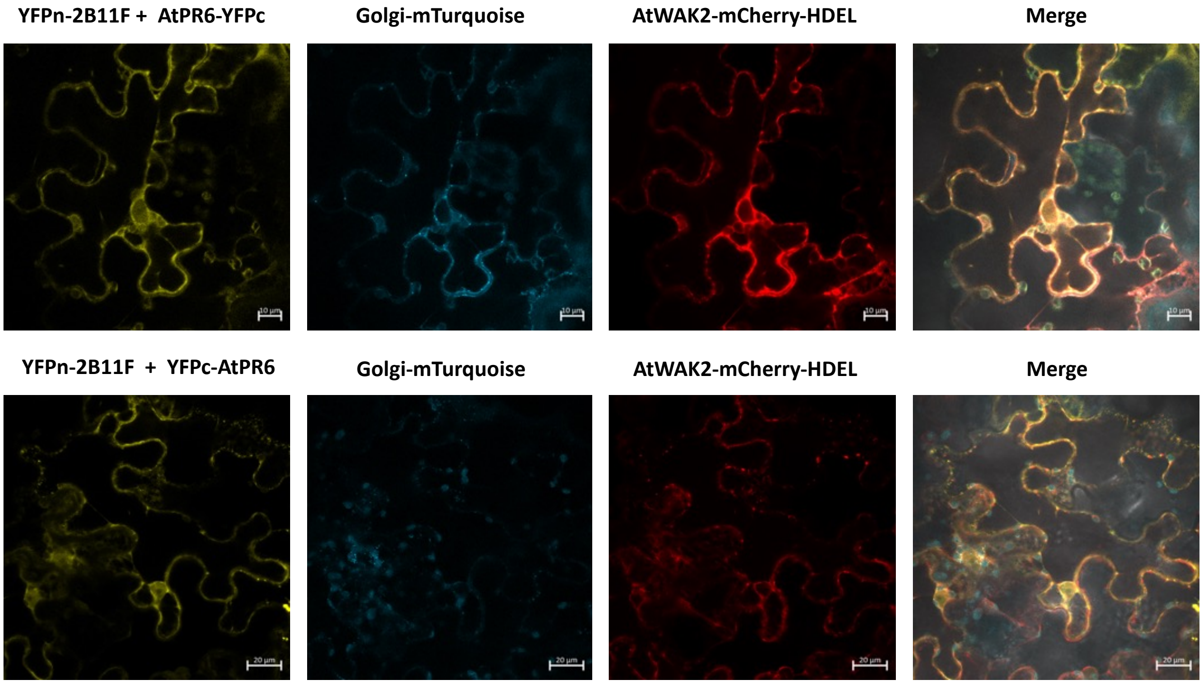


**Fig S4**. *Hs*2B11 and *At*PR-6 interact in the ER in *N. benthamiana* epidermal cells. Co-localization of YFPn-*Hs*2B11 and *At*PR-6-YFPc or YFPc-*At*PR-6 and the ER marker *At*WAK2-mCherrry-HDEL. Bars=10 or 20 μm as indicated in the figures. Cyan fluorescent protein (CFP) fused to a Golgi signal peptide is used to confirm expression of the T-DNA insert.

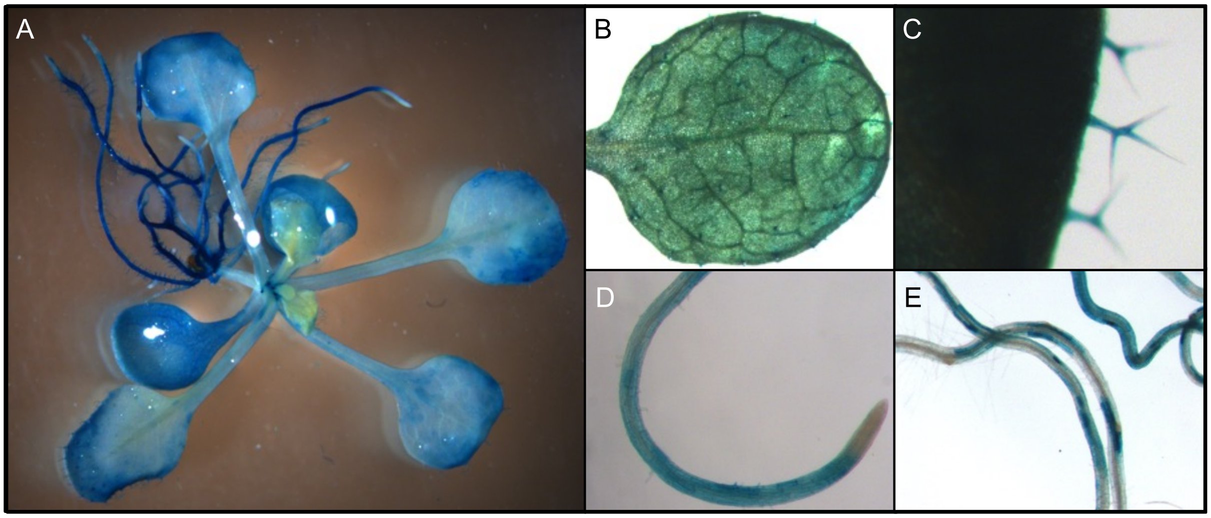


**Fig S5.** Histochemical localization of GUS activity controlled by the *AtPR6* promoter in Arabidopsis after a 4 h incubation with X-Gluc at 37°C. GUS activity controlled by *AtPR6* promoter in uninfected whole (A) Arabidopsis seedlings, (B) leaves (C) emerging leaves with trichomes, and (D-E) roots.

**Table S1**. Y2H summary using *At*PR-6 as a bait.

| Locus | Function | |
|---|---|---|
| AT1G20160. | ATSBT5.2, CO2 RESPONSE SECRETED PROTEASE, CRSP, SBT5.2 | **Proteases** |
| AT3G14240 | Subtilase family protein | |
| AT3G14240 | Subtilase family protein | |
| AT3G52500 | Eukaryotic aspartyl protease family protein | |
| AT1G10810 | NAD(P)-linked oxidoreductase superfamily protein | **Redox** |
| AT1G10810 | NAD(P)-linked oxidoreductase superfamily protein | |
| AT1G10810 | NAD(P)-linked oxidoreductase superfamily protein | |
| Hsc_gene_13411 | have a signal peptide = unknown | **Nematodes genes** |
| Hsc_gene_13411 | have a signal peptide = unknown | |
| HS_G3197 | Unknown | |
| Hsc_gene_20080 | Transthyretin-like with SP | |
| AT3G57520 | ATSIP2, RAFFINOSE SYNTHASE 2, RS2, SEED IMBIBITION 2, SIP2 | **Metabolism** |
| AT3G57520 | ATSIP2, RAFFINOSE SYNTHASE 2, RS2, SEED IMBIBITION 2, SIP2 | |
| AT5G20250 | DARK INDUCIBLE 10, DIN10, RAFFINOSE SYNTHASE 6, RS6 | |
| AT2G36530 | ENO2, ENOLASE 2, LOS2, LOW EXPRESSION OF OSMOTICALLY RESPONSIVE GENES 2 | |
| AT2G36530 | ENO2, ENOLASE 2, LOS2, LOW EXPRESSION OF OSMOTICALLY RESPONSIVE GENES 2 | |
| AT2G22125 | CSI1, POM2 \| POM-POM 2, cellulose synthase. | **CAZymes** |
| AT1G78060 | Glycosyl hydrolase family protein | |
| AT1G68560 | ALPHA-XYLOSIDASE 1, ALTERED XYLOGLUCAN 3, ATXYL1, AXY3, GH31, THERMOINHIBITION RESISTANT GERMINATION 1, TRG1, XYL1 | |
| AT3G23640 | Symbols: HGL1 \| heteroglycan glucosidase 1 | |
| AT1G19220 | ARF11, ARF19, AUXIN RESPONSE FACTOR 19, AUXIN RESPONSE FACTOR11, IAA22, INDOLE-3-ACETIC ACID INDUCIBLE 22 | **Diverse** |
| AT1G19220 | ARF11, ARF19, AUXIN RESPONSE FACTOR 19, AUXIN RESPONSE FACTOR11, IAA22, INDOLE-3-ACETIC ACID INDUCIBLE 22 | |
| AT1G04870 | ATPRMT10, PRMT10, PROTEIN ARGININE METHYLTRANSFERASE 10 | |
| AT5G37740 | Calcium-dependent lipid-binding (CaLB domain) family protein;(so | |
| AT1G70730 | PGM2 \| phosphoglucomutase 2 | |
| AT5G57630 | SnRK3.4, CIPK21 \| SNF1-RELATED PROTEIN KIN | |
| AT4G00680 | Symbols: ADF8 \| actin depolymerizing factor 8 | |
| AT5G23820.1 | MD2-RELATED LIPID RECOGNITION 3, ML3 | |
| AT4G02060 | MCM7, PRL, PROLIFERA | |
| AT3G61440 | ARATH;BSAS3;1, ATCYSC1, BETA-SUBSTITUTED ALA SYNTHASE 3;1, CAS-C1, CYSC1, CYSTEINE SYNTHASE C1, ß-CYANOALANINE SYNTHASE C1 | |
| AT3G29680 | HXXXD-type acyl-transferase family protein | |
| AT5G07470 | ARABIDOPSIS THALIANA METHIONINE SULFOXIDE REDUCTASE 3, ATMSRA3, PEPTIDEMETHIONINE SULFOXIDE REDUCTASE 3, PMSR3 | |

**Table S2.** Primers used in this study.

| Purpose | Primers name | Sequence |
|---|---|---|
| ***GATEWAY Cloning*** | 2B11_ATG_F | AAAAAGCAGGCTTCACCATGGAGGAATTGTTTCTTTA |
| | 2B11_STOP_R | AGAAAGCTGGGTGTCATAAATTAATTCTTCTTCCACCGTAAT |
| | 2B11_noSTOP_R | AGAAAGCTGGGTGTAAATTAATTCTTCTTCCACCGTAAT |
| | Nterm_2B11_noSTOP_R | AGAAAGCTGGGTGTTGGTCGATATATAAATTTCCATTG |
| | Nterm_2B11_STOP_R | AGAAAGCTGGGTGTCATTGGTCGATATATAAATTTCCATTG |
| | Cterm_2B11_ATG_F | AAAAAGCAGGCTTCACCATGTCGTCAGATGGGTGTTCTCGC |
| | 2B11m_ATG_F | AAAAAGCAGGCTTCACCATGGACTGTGGTGGATCTTTGCG |
| | 2B11m_STOP_R | AGAAAGCTGGGTGTCATAACTTAAGATTTCGCGCTATTGA |
| | 2B11m_noSTOP_R | AGAAAGCTGGGTGTAACTTAAGATTTCGCGCTATTGA |
| | AtPR6_ATG_F | AAAAAGCAGGCTTC ACCATGTCGACCGAATGTCCTAGGAAGA |
| | AtPR6_STOP_R | AGAAAGCTGGGTGCTAGCCGGATTTGGGGGTTTTGACGA |
| | AtPR6_noSTOP_R | AGAAAGCTGGGTGGCCGGATTTGGGGGTTTTGACGA |
| | AttB1 adapter | GGGGACAAGTTTGTACAAAAAAGCAGGCT |
| | AttB2 adapter | GGGGACCACTTTGTACAAGAAAGCTGGGT |
| **Transgenic plant** | PR6OE_F_BamH1 | tataggatccATGTCGACCGAATGTCCTAGGAAGA |
| | PR6OE_R_Sst1 | TATAGAGCTCCTAGCCGGATTTGGGGGTTTTGACGA |
| | Hs2B11OE_F_BamH1 | TATAGGATCCATGGGAGGAATTGTTTCTTTACTATC |
| | Hs2B11OE_R_Sst1 | tatagagctcTCATAAATTAATTCTTCTTCCACC |
| | PR6Pro_F_Xba1 | GAGATCTAGAGGCAGTGTGAAACATACACAAAACTAAACAC |
| | PR6Pro_R_BamH1 | tctcggatccgtagtcttctgatgtatttgtatgctc |
| **Y2H** | 2B11_Bait_F_EcoR1 | tatagaattcGGAGGAATTGTTTCTTTACTATCAAGAAG |
| | 2B11_bait_R_BamH1 | tgtgggatccTCATAAATTAATTCTTCTTCCACCG |
| | AtPR-6_bait/prey_F_EcoR1 | tatagaattcATGTCGACCGAATGTCCTAGGAAGA |
| | AtPR-6_bait/prey_R_BamH1 | TATAggatccCTAGCCGGATTTGGGGGTTTTGACGA |
| | Hs2B11_Nterm_Bait_R_BamH1 | GCGCGGATCCTCATTGGTCGATATATAAATTTCCA |
| | Hs2B11_Cterm_F_EcoR1 | TATAGAATTCTCGTCAGATGGGTGTTCTCGCATA |
| | Hs2B11_3R/1R_F_EcoR1 | tatagaattcGACTGTGGTGGATCTTTGCGTATT |
| | Hs2B11_M_R_BamH1 | GCTAggatccTCAGTCTAACTTAAGATTTCGCGC |
| **Genotyping KO** | LP111051C | TTGTCTACACGTTGCCACTTG |
| | RP111051C | TGGTCATTTTTGTCGGCTAAG |
| | LBB1.3Plus | ATTTTGCCGATTTCGGAAC |
| **BiFC** | 2B11FullF2 | ccatggctATGGGAGGAATTGTTTCTTTACTATCA |
| | 2B11FullR2 | actagtTAAATTAATTCTTCTTCCACCGTAATCAATTC |
| | 2B11-Cterm_F | ccatggctATGTCGTCAGATGGGTGTTCTCGCA |
| | PR6_F | GGGTCCCTTTCGACCGAATGTCCTAGGAAGAACTCGTG |
| | PR6_R | TATCCGGAGCCGGATTTGGGGGTTTTGACGACGATACGGT |
| **Sanger Sequencing** | 2BFInR | CCAAAATTATTTTCAGCACCATT |
| | 2bFInF | CAAATGGAAAGCAGTCAGATTG |
| | P35S | gatgacgcacaatcccactatc |
| | T35Sm | ctccagaataatgtgtgagtag |
| | U | TGTAAAACGACGGCCAGT |
| | R-1 | CAGGAAACAGCTATGACC |
| **qPCR** | 2B11F_qPCR_F | CAAGAAGACAAGCACTAAAGCA |
| | 2B11F_qPCR_R | TCCCAATTCACACTCAAATCC |
| | AtPR6_qPCR_F | GGAACAAATGGTGACTATGCG |
| | AtPR6_qPCR_R | TGACGACGATACGGTTTCCA |
| | qUBOX_F | TGCGCTGCCAGATAATACACTATT |
| | qUBOX_R | TGCTGCCCAACATCAGGTT |
